# Systematic assessment of lipid profiles for the discovery of tissue contributors to the circulating lipid pool in cold exposure

**DOI:** 10.1101/2021.11.12.468392

**Authors:** Raghav Jain, Gina Wade, Irene Ong, Bhagirath Chaurasia, Judith Simcox

**Author notes:** Corresponding Author: Judith Simcox.

## Abstract

Plasma lipid levels are altered in chronic conditions such as type 2 diabetes and cardiovascular disease as well as acute stresses such as fasting and cold exposure. Advances in mass spectrometry based lipidomics have uncovered the complexity of the plasma lipidome which includes over 500 lipids that serve functional roles including energy substrate and signaling molecule. The plasma lipid pool is maintained through regulation of tissue production, secretion, and uptake. A major challenge is establishing the tissues of origin and uptake for various plasma lipids, which is necessary to determine the lipid function. Using cold exposure as an acute stress, we performed global lipidomics on the plasma and nine tissues that may contribute to the circulating pool. We found that numerous species of plasma acylcarnitines (ACars) and ceramides were significantly changed with cold exposure. Through computational assessment, we identified the liver and brown adipose tissue (BAT) as major contributors and consumers of circulating ACars, in agreement with our previous work. We further identified the kidney and intestine as novel contributors to the circulating ACar pool and validated these findings with gene expression analysis. Regression analysis also identified that the BAT and kidney as regulators of the plasma ceramide pool. These studies provide an adaptable computational tool to assess tissue contribution to the plasma lipid pool. Our findings have implications in understanding the function of plasma ACars and ceramides, which are elevated in metabolic diseases.

**Summary:** There are over 500 identified lipids in circulating plasma, many without known origin or function. Using untargeted lipidomics on plasma and nine other tissues of cold exposed mice, we identified novel regulation of circulating acylcarnitines through the kidney and intestine, and a multiorgan system that regulates plasma ceramides. Our findings offer new targets for the study and functional characterization of circulating lipids in acute cold exposure and a computational resource for other investigators to explore multi-tissue lipidome remodeling during cold exposure.

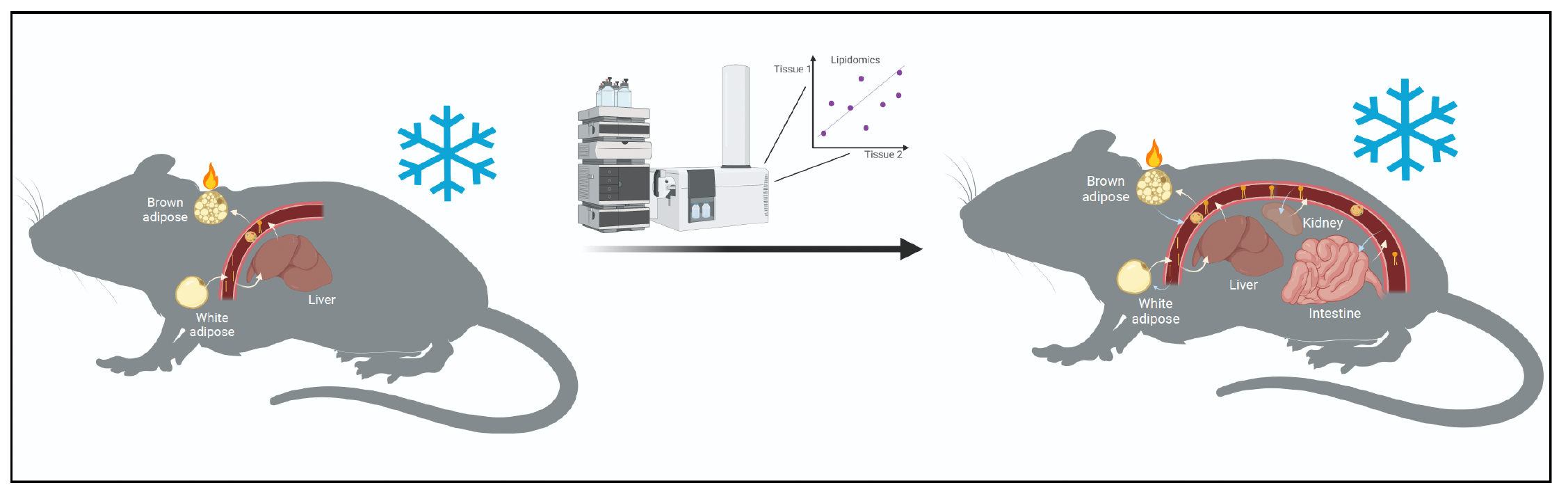

**Highlights:**

- Global lipidomics atlas of 9 tissues and plasma demonstrate dynamic shift with cold exposure.
- Adaptive resource for the selection of extraction method, data processing, and data analysis of multi-tissue global lipidomics data.
- Regression analysis identified the liver, BAT, intestine, and kidney as regulators of the plasma acylcarnitine pool that are not apparent by lipid levels alone.
- Acute cold exposure increases plasma ceramide levels, with the BAT and kidney as major contributors

## Introduction

Plasma lipids reflect metabolic stress and disease. The advent of mass spectrometry-based lipidomics has expanded the use of plasma lipids as diagnostic markers including newborn screens for inborn errors of metabolism to assess specific plasma acylcarnitines (ACars) (Fletcher, 2008; Yoon, 2015). These technological advances have also determined that there are more than 500 identified lipids in the human plasma lipidome (Quehenberger and Dennis, 2011). These lipids have diverse structure with variations in head group, backbone, and acyl chain configuration. Despite the extended knowledge that plasma lipids can indicate metabolic abnormalities and disease, little is known about the functional role, regulation, or tissue of origin for the majority of lipids found in the circulation.

The abundance of plasma lipids is dynamically regulated to respond to nutrient availability. In fasting, there is an increase in plasma free fatty acids (FFAs) which is driven by white adipose tissue (WAT) lipolysis, signaling a shift from glucose to lipid catabolism. In the fed state, a rise in dietary triglycerides, packed in chylomicrons, coincides with glucose stimulated insulin secretion to induce adipocyte lipid storage. Beyond nutrient availability, factors such as age, diet, sex, and genetic background regulate lipid uptake and secretion into plasma, demonstrating the tight regulation of the plasma lipidome (Linke et al., 2020; Surma et al., 2021). In addition to serving as energy substrates, plasma lipids have been recognized as endocrine signals with novel classes of lipids such as fatty acid esters of hydroxy fatty acids (FAHFAs) regulating insulin sensitivity (Yore et al., 2014). More work is needed to understand how the physiological context affects the complex fates of plasma lipids.

Efforts to determine the regulation and function of lipids and lipidome remodeling have utilized acute stresses such as fasting, circadian rhythm disruption, and cold exposure to rapidly alter tissue lipid profiles (Banfi et al., 2018; BasuRay et al., 2017; Simcox et al., 2017; Xia et al., 2021). Cold exposure is an energy demanding, selective pressure that requires rapid mobilization of lipids as a fuel and signal to activate mitochondrial oxidation (Park et al., 2019). By using the stress of cold exposure, interorgan lipid regulatory pathways have been identified including the regulation of adipocyte lipolysis by hepatic insulin signaling, and the regulation of bile acid production and secretion (Sostre-Colón et al., 2021; Worthmann et al., 2017). Our previous work utilized cold exposure to determine that plasma ACars are produced through multi-tissue lipid processing. In this system, cold activates the release of FFAs from the WAT for uptake by the liver. This FFA uptake results in transcriptional activation of liver ACar production and export into the plasma, where ACars are taken up by the brown adipose tissue (BAT) to serve as a fuel source for thermogenesis (Simcox et al., 2017). These studies were the first to identify and track the complexity of lipid processing through multiple tissues, and to characterize a functional role for ACars in thermogenesis. The results further highlight the utility of cold exposure to delineate the regulation and function of plasma lipids.

We built upon our previous work to better understand the contribution of various tissues to the plasma lipidome in acute cold exposure. After optimizing the lipid extraction for a broad range of tissues, we performed global lipidomics on the plasma, liver, BAT, inguinal WAT (iWAT), epidydimal WAT (eWAT), kidney, intestine, lung, heart, and gastrocnemius skeletal muscle (GSM) of mice kept at room temperature or exposed to cold (4°C) for 6h. We observed the greatest cold-induced lipid changes in the liver, BAT, and plasma, with the most dynamic lipid classes being ACars, triglycerides, and sphingolipids. Through correlation analysis, we were able to replicate previous observations that ACars from the plasma are produced in the liver and go to the BAT for catabolism. Regression analysis identified previously unknown regulation of these plasma ACars by the intestine and kidney. Gene expression analysis confirmed a likely role for the intestine as a tissue of uptake and the kidney as a site of production. We extended this analysis and found that ceramides (Cers), a type of signaling sphingolipid, are increased in the plasma with acute cold exposure. Regression analysis showed that the BAT is a major regulator of the plasma Cer pool. These studies demonstrate that numerous tissues contribute to the circulating lipid pool of ACars and Cers and determined that computational assessment was able to identify novel tissue contribution. Future work will be needed to establish the functional role of plasma Cer species in acute cold exposure.

## Results

### Optimization of lipid extraction methodology is required for multi-tissue analysis

The method of tissue homogenization and organic extraction affects lipid recovery (Reis et al., 2013). To evaluate the impact of extraction method on lipid recovery in various tissues, we tested four different solvent extractions including Folch, methyl-tertbutyl-ether (MTBE), acidified MTBE (acidic), and isopropanol (IPA). The Folch, MTBE, and acidic methods allow lipid recovery by separation of an aqueous and lipid-containing organic phase (**Figure 1A**). In contrast, the single-phase IPA method relies on precipitation of a non-recoverable fraction (NR) with lipids suspended in solution. We pooled lipid extracts for plasma from three mice and performed LC-MS/MS analysis to determine total spectral features and lipid identifications in positive and negative ionizations using LipidAnnotator. Computational filtering was applied to ensure unique and high confidence lipid annotation. The Folch method yielded the highest number of total spectral features (3912), but fewest lipid identifications (278 lipids) of all methods. The MTBE method captured the highest number of identifications (368 lipids) for plasma (**Figure 1B**). Because lipid extractions perform differently based on tissue type (Höring et al., 2021), we repeated the extractions for the GSM and liver (**Figure S1A**). The Folch method resulted in the highest number of features, but the IPA extraction provided the most lipid identifications for both liver (638) and GSM (434). These results indicated that the IPA method performed well for solid tissue extractions, but the MTBE method had slightly higher lipid annotation for plasma.

**Figure 1:**
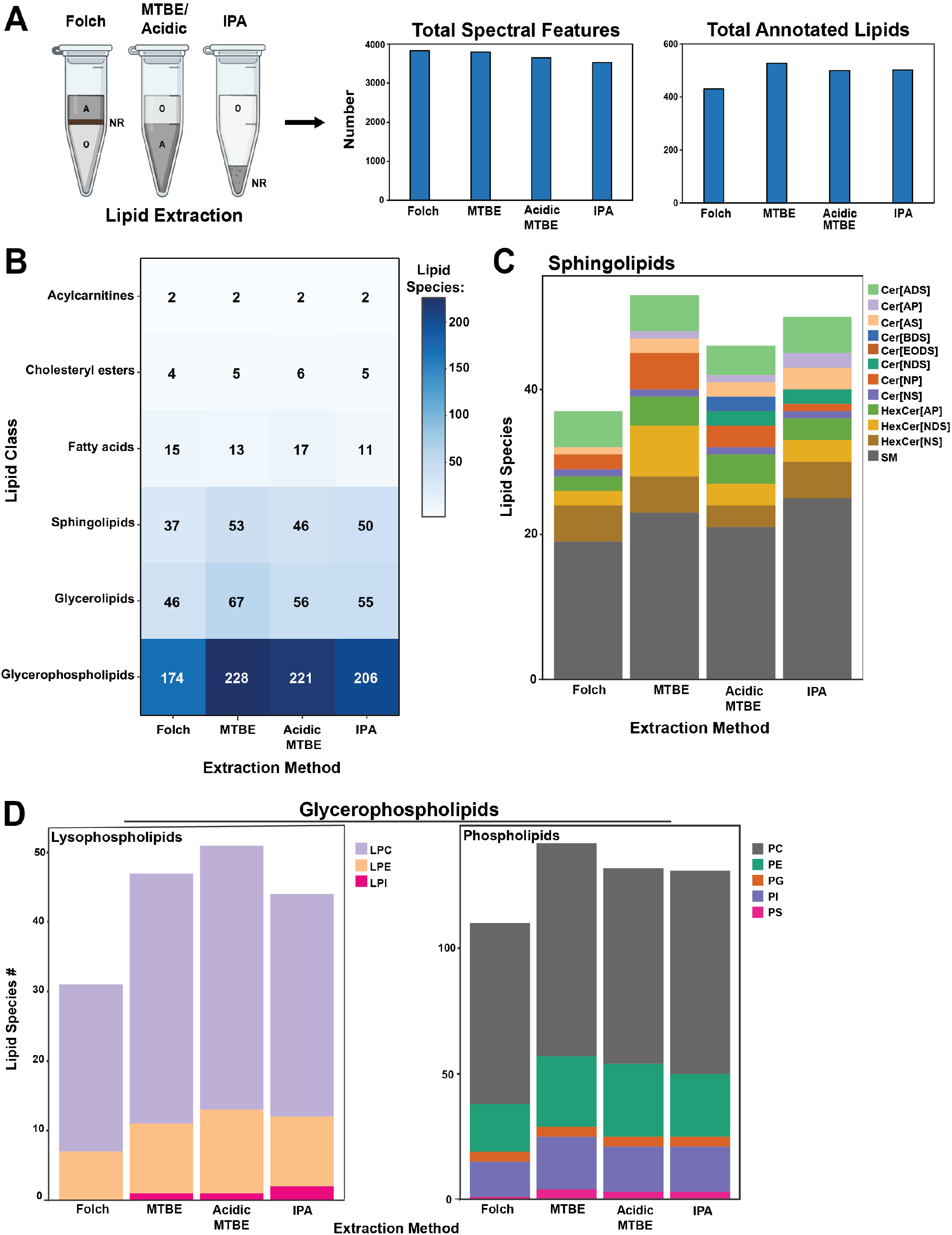
Extraction methods enrich for different lipid classes. A) Comparison of phase separation between two-phase Folch and MTBE, and single phase IPA extractions where A = aqueous, O = organic, NR = non-recoverable (protein). Total spectral features, and total lipid identifications shown. Lipids were identified in LipidAnnotator following LC/QTOF-MS/MS data collection on pooled plasma from male C57BL6J mice (n=3). B) Heat map showing the distribution of lipid identification between the four extractions. C) Sphingolipid identification between the four extractions with distribution across subclasses shown by bar color. D) Distribution of glycerophospholipids split by lysophospholipids and phospholipids between the four methods. Bar color indicates further breakdown into subclasses. Data represents summed features and annotations from positive and negative ionization modes after manual curation.

To determine if differences in identified lipids were driven by overall lipid recovery or due to preferential extraction of lipids between methods, we compared the number of identifications for each lipid class across the extractions. In plasma, the Folch method had the lowest identifications of all the main lipid classes, including sphingolipids, glycerolipids, and glycerophospholipids (**Figure 1B**). In contrast, the MTBE method had the most identifications, but was only slightly higher than the IPA method with ~40 unique lipids varying between the groups; the major contributing lipid class to these differences was glycerophospholipids. We further delved into these differences by looking at the breakdown of lipid subclasses within each major class. The greatest variance was observed in sphingolipids. We noticed that although the total number of sphingolipids was similar between MTBE and IPA extractions, the distribution of sub-classes indicated greater hexosylceramide (HexCer) recovery in MTBE (**Figure 1C**). This was offset by more sphingomyelin species in the IPA extraction, as well as the identification of Cer[NDS] (ceramide non-hydroxy fatty acid dihydrosphingosine) species which were not detected in MTBE. Phosphatidylcholine species, the most abundant phospholipid in mammals, were the main lipids enriched in MTBE compared to IPA extraction (**Figure 1D**). No lysophosphatidylinositols were detected in the Folch extractions and there was a general decrease in lysophospholipids detected in Folch compared to all other methods. These findings are relevant in the context of targeted studies in plasma, which may prioritize the detection of certain lipids over others.

The IPA method had consistent and comprehensive annotation for all lipid classes and subclasses when compared to the other extractions in the liver (**Figure S1**). This was also true in GSM, except for sphingolipids, which were enriched in the Folch method compared to all others (**Figure S1**). In GSM, the increased identification of sphingolipids in the Folch method was due to higher detection of ceramide species (>27-32 species) but lower sphingomyelin (<4-7 species). Interestingly, we saw nearly double the number of lysophospholipid species in acidic than IPA, Folch, or MTBE methods in GSM but not in liver or plasma. This is potentially caused by increased hydrolysis of PC and PE species, which are highly enriched in GSM, since acidic environments can promote phospholipid breakdown (Zuidam and Crommelin, 1995). We concluded that the IPA method was best suited for the extraction of different tissue and the recovery of diverse lipid species.

### Individual tissues have distinct cold induced lipid remodeling

Acute cold exposure is a metabolic stress that induces rapid lipid remodeling across multiple tissue (Park et al., 2019; Von Bank et al., 2021). This lipid remodeling is driven by increased adipose tissue lipolysis which then leads to hepatic steatosis and elevated plasma FFAs and TGs (Heine et al., 2018; Mottillo et al., 2014; Sostre-Colón et al., 2021). These lipolysis induced changes are similar to phenotypes of fasted and high fat diet fed mice but occur on a faster time scale with major changes in 5 hours of cold exposure, mimicking 24 hours of fasting (Guan et al., 2009; Heine et al., 2018; Inagaki et al., 2007; Perry et al., 2014; Simcox et al., 2017). We have previously employed the model of cold exposure to demonstrate a functional role for plasma ACars as fuel for BAT thermogenesis (Simcox et al., 2017). This finding highlights the plasticity of lipids in response to physiologic stress and the potential of rapid, cold-induced lipid remodeling to provide insight into lipid functions. The observations also unearthed the complexity of lipid processing contributing to the plasma lipid pool since ACar production is induced by WAT lipolysis of triglycerides into FFAs, which are then processed into ACars by the liver, secreted into circulation, and taken up by BAT (Simcox et al., 2017). These studies were limited by a focused exploration of known contributors to the plasma ACar pool. More work is needed to understand the tissue of origin for other circulating lipid as well as the functional contribution of other tissues to thermogenesis.

To identify tissues of origin and processing for the plasma lipid pool, we placed mice in cold (4°C) or control room temperature (RT, 24°C) for 6 hours while fasting, then harvested ten tissues for LC-MS lipidomics. Data was normalized to individual internal standards for each lipid class and starting tissue amount, and was reported as semi-quantitatively in either nM lipid for plasma or nmol lipid/g for all other tissue. There were 166 lipids commonly detected across all nine solid tissue, and principal component analysis (PCA) analysis based on these lipids showed clustering by tissue type in RT and cold (**Figure 2A**). There was a noticeable shift in clustering in cold exposed mice, but the tissue overlap pattern remained similar. Regardless of temperature, BAT did not overlap with any other tissue whereas iWAT and eWAT, the main lipid storage depots, clustered together but not with other tissue. For the remaining tissue, there was a high degree of overlap in RT, while distinct clustering emerged for GSM and heart in cold exposure.

**Figure 2:**
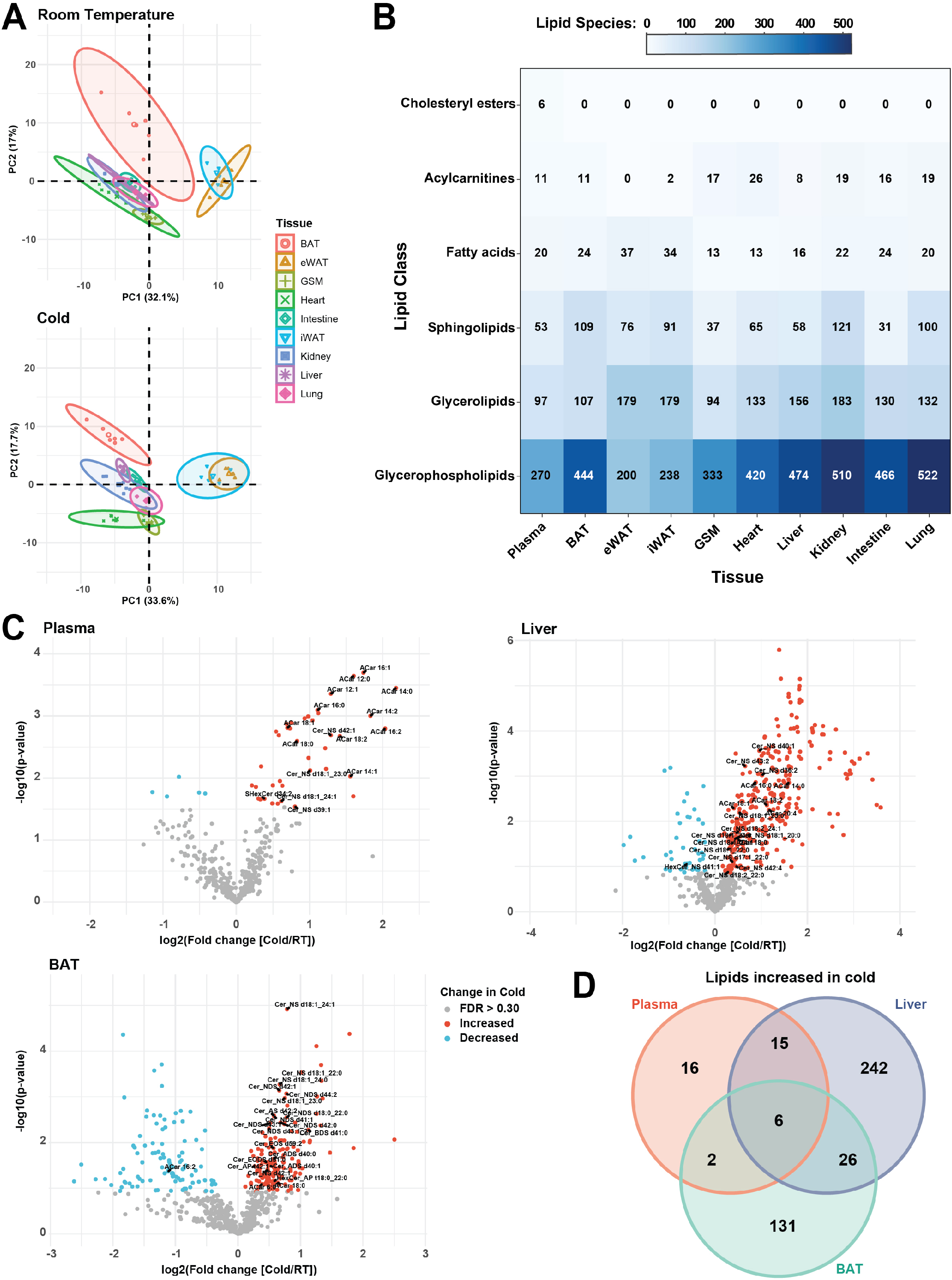
Tissue lipid composition undergoes rapid and dynamic remodeling with cold exposure. A) Principal component analysis between 9 tissues in room temperature (RT, 24°C) and cold exposure (Cold, 4°C). B) Heat map showing lipid class distribution between 9 tissues and plasma extracted by IPA method. C) Volcano plots illustrating significant and high fold change between RT and Cold in plasma, liver, and brown adipose tissue (BAT). D) Venn Diagram showing overlap of lipid species significantly increased between RT and Cold in the plasma, liver, and BAT (n=6 per condition). Student’s t-test was used to determine significance in volcano plots with adjustment for multiple comparisons. FDR *q*<0.30 red dots indicate significant increases in cold and blue is significant decreases in cold.

Next, we looked at the distribution of individual lipid species detected in each tissue for different lipid classes (**Figure 2B**). We detected the fewest number of total lipids in plasma (457) and the most in kidney (855). The majority of identified lipids were glycerophospholipids though unsurprisingly, a significant proportion of the WAT depots were composed of glycerolipids. Interestingly, in BAT, we saw more sphingolipid species (109) than glycerolipids (107) which was not observed in any other tissue. There were modest trends in lipid abundances when comparing lipid classes in RT and cold for the various tissues (**Figure S2A**). There was a significant increase in liver triglycerides and FFAs, reflecting the hepatic steatosis induced by cold exposure (Sostre-Colón et al., 2021). In both WAT depots, there was an increase in FFAs and diacylglycerols, consistent with increased adipose lipolysis (Heine et al., 2018). Interestingly, we did not see a significant decrease in WAT triglycerides associated with lipolysis, which could be due to the normalization by tissue weight rather than fat mass.

To gain insight into which individual lipid species contributed to cold-induced lipid remodeling, we compared the number of lipids significantly changed due to housing temperature after false discovery correction (*q*<0.30) in each tissue (**Figure 2C and Figure S2B; Figure S1A**). We found that lung, intestine, and kidney – the tissue with the most identified lipids – had very few significant changes in lipids during the shift from RT to cold (**Figure S2B**). Plasma and liver generally had more lipids increased in cold, driven by increased ACar and Cer species in plasma (**Figure 2C**) and increased TG and Cer species in the liver. The most dynamic lipid changes occurred in BAT as there were 165 lipids significantly increased and 107 lipids decreased in cold. The majority of lipids decreased in cold BAT were triglycerides (82 species) which coincides with increased lipolysis to fuel thermogenesis (Sveidahl Johansen et al., 2021). Unlike plasma or liver, the main lipids increased in cold exposed BAT were cardiolipins and phosphatidylcholine species. Cardiolipins are important lipids for the inner mitochondrial membrane and their increase is due to higher mitochondrial content in BAT to support thermogenesis (Lynes et al., 2018; Sustarsic et al., 2018). There were also significant increases in Cers and ACars in cold BAT.

Because the highest magnitude changes in lipid composition with cold exposure occurred in the plasma, liver, and BAT, we compared the cold induced lipid changes in the tissue. There were only six lipids commonly increased in all three tissues – ACar 18:0, Cer_NS d18:1_23:0, Cer_NS d18:1_24:1, FA 20:3, FA 22:4 and PC 34:4 (**Figure 2D; STable 1**). There was also more overlap in commonly increased lipid species between plasma and liver (15 lipids) than between plasma and BAT (2 lipids). Together these data indicated a unique lipid profile for each tissue with a subset of tissues exhibiting cold induced shifts in lipid composition. In particular, the plasma, liver, and BAT had lipid remodeling that was largely distinct from each other but shared some features.

### Tissue of origin and processing for acylcarnitines can be predicted using correlation analysis

One of the shared lipids increased in cold exposure between plasma, liver and BAT was ACar 18:0. Due to the established relationship of circulating ACars originating from liver and fueling BAT, we compared the most abundant ACar species detected in each tissue during RT and cold (Simcox et al., 2017). All major ACar species in plasma and most in liver were increased during cold exposure (**Figure 3A**). In the plasma, there were no detected ACars with an acyl chain greater than 18 carbons, and no detected ACars with acyl chains below 14 carbons. This suggested a greater export of medium (<14 carbon) ACars from the liver when compared to long chain ACars. In BAT, only ACar 18:0 was increased with cold exposure, but this increase was not significant (**Figure 3A**).

**Figure 3:**
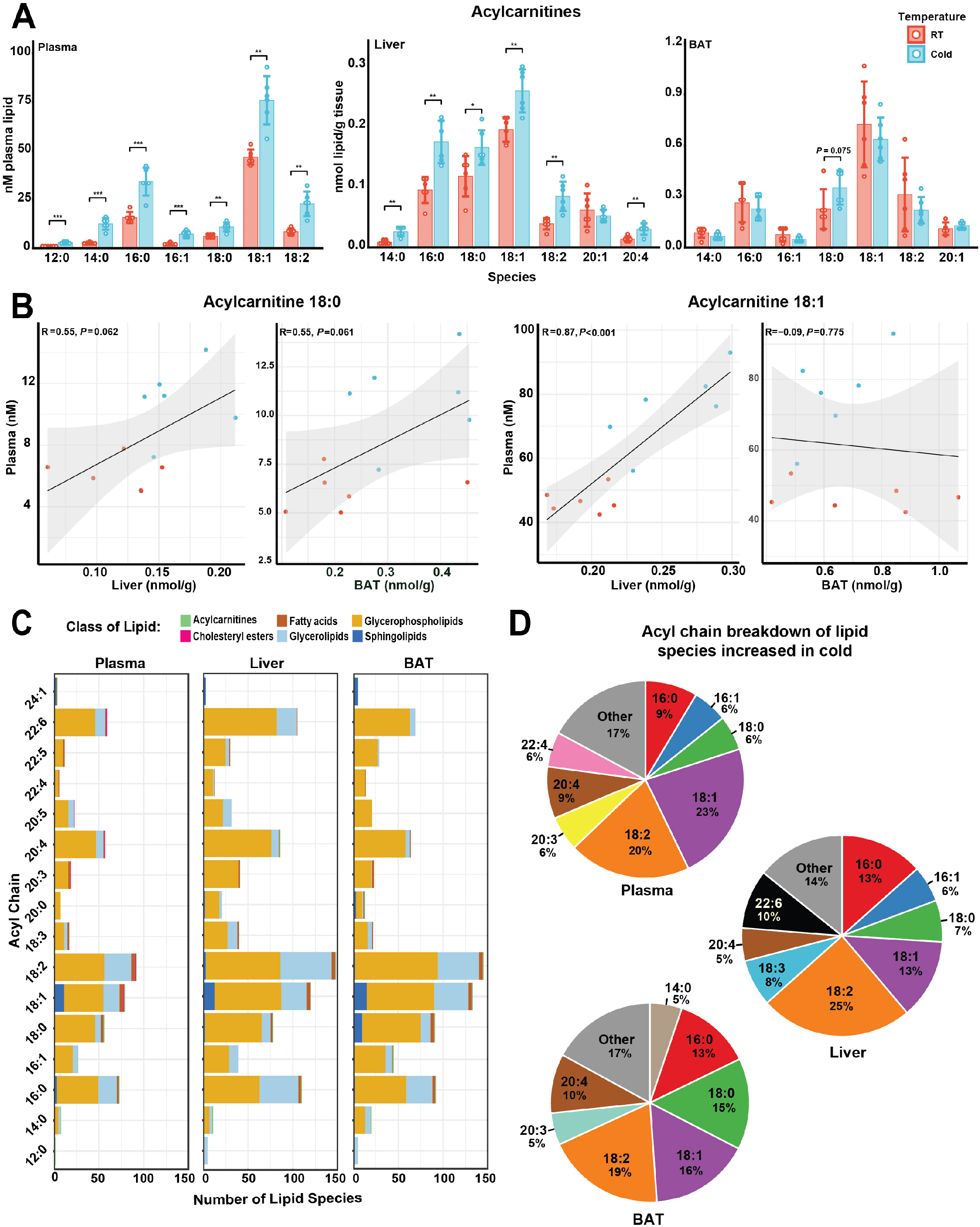
Correlation analysis reveals a plasma, liver, BAT axis for acylcarnitine processing. A) Acylcarnitine (ACar) species abundance in plasma, liver, and BAT between room temperature (RT, 24°C) and cold exposure (Cold, 4°C). B) Correlation analysis between plasma, liver and BAT for ACar 18:0 and ACar 18:1. C) Acyl chain distribution showing the prevalence of different chains across the plasma, liver, and BAT lipidomes. Lipid class occurrence of a given acyl chain is indicated by color. D) Percentage acyl chain composition of lipids increased in cold from Fig 2C. Only acyl chains occurring in at least 5% of increased lipids are shown. Student’s t-test used for all pairwise comparisons; \**P*<0.05 \*\**P*<0.01 \*\*\**P*<0.001. (n=6)

To measure the relationship of ACars between the plasma, liver and BAT, we correlated ACar 18:0, the only ACar increased in all three tissue, and ACar 18:1, the most abundant ACar in all three tissues (**Figure 3B**). Plasma ACar 18:0 was positively correlated with both liver and BAT with an identical *R* = 0.55, and *P* = 0.062 and 0.061, respectively. These data were indicative of a general increase in ACar 18:0 in liver that caused an increase in plasma ACar 18:0 and deposited in BAT for thermogenesis (Simcox et al., 2017). In contrast, ACar 18:1 was strongly correlated between plasma and liver (*R* = 0.87, *P*<0.001) but shared no correlation between plasma and BAT (*R* = −0.09, *P* = 0.775). The absence of correlation was similar across other ACars detected in all three tissues (**Figure S3A**). Upon closer inspection, the lack of significant correlation between plasma and BAT ACars seemed to be driven by RT levels (red dots), whereas in cold, BAT ACars trended towards a positive correlation with plasma levels (blue dots).

Our findings recapitulated previously published results demonstrating that cold exposure induces hepatic production of ACars that are secreted into the plasma to fuel thermogenesis in BAT (Simcox et al., 2017). Cold-induced catabolism of ACars would prevent accumulation in BAT. These data suggested that cold exposure could be used to computationally infer the individual lipid contribution of different tissue to the plasma pool. The data also demonstrated acyl chain variation in ACar species between each tissue. We next assessed acyl chain abundance in the various tissues to determine if presence, or absence, of specific acyl chains during cold exposure could distinguish between tissues. This information could be used to track the lipid contribution of various solid tissue to the plasma lipid pool.

### Tissue-specific acyl chain signatures are maintained in cold exposure

Under steady state conditions, each tissue has a distinct acyl chain composition to each lipid class (Harayama and Riezman, 2018). To determine whether tissue specific acyl chain signatures are maintained in cold exposure, we first compared the number of lipid species containing a given acyl chain in plasma, liver, and BAT (**Fig 3C**). General acyl chain distribution patterns were largely similar across all three tissues. The most common acyl chains were 18:2 (first) and 18:1 (second) and existed predominantly in phospholipids, in agreement with previous studies (Harayama and Riezman, 2018; Oemer et al., 2020). The third most common acyl chain was 16:0 with 73 lipids in the plasma, 110 lipids in the liver, and 92 lipids in the BAT containing this acyl chain. In BAT, 18:0 containing lipids were also common (91 lipids). Upon further inspection, we noticed that this increase in 18:0 containing species was driven by sphingolipids (10 sphingolipids in BAT), which was more than in plasma (1 sphingolipid) or liver (2 sphingolipids). We next looked at the total intensity of lipids containing various acyl chains and observed the greatest percentage shift in liver and BAT, while the total proportions of acyl chain composition remained similar, suggesting that individual lipids species are driving the cold-induced changes (**Figure S3B**).

Next, we compared the major acyl chain distribution (>5%) of lipids increased in cold for plasma, liver, and BAT (**Figure 3D**). Similar to general acyl chain prevalence, 18:2 was the most common acyl chain in lipids increased across all three tissues, and 18:1 was second. In BAT only, 15% of the acyl chain composition of lipids increased in cold contained 18:0, whereas this number was less than 10% for plasma and liver. These data indicated that an acyl chain signature may differentiate BAT from other tissue. In particular, the presence of sphingolipids containing the 18:0 carbon chain was increased compared to plasma or liver. The 18:0 containing sphingolipids present in BAT were from ceramides, and the 18:0 was specifically from the sphingoid base (**Figure S3C**). These ceramides were also detected in iWAT and eWAT but not plasma or liver. Total 18:0 ceramides were not increased during cold in BAT, but they were significantly increased in iWAT and eWAT. Therefore, 18:0 containing ceramides may be lipid signatures for BAT and WAT, an observation that aligns with their known regulation of beige adipocyte differentiation (Chaurasia et al., 2016; Chaurasia et al., 2021). Since these ceramides were not detected in the plasma, their functions may be intra-tissue rather than related to inter-tissue signaling.

### Regression analysis identifies novel tissue contributions to the plasma acylcarnitine pool

We sought to extend our ACar observations and take an unbiased, lipidome-wide approach to potentially identify unknown contributors of the plasma ACar pool. We regressed each lipid identified in our analysis against the most abundant plasma ACars to generate a list of tissue lipids that were significantly predictive of plasma ACars (**Figure 4A**). Distinct patterns of lipids from a given tissue were either positive or negative predictors of plasma ACars. Liver and iWAT lipids were generally positively predictive of plasma ACars which supports the known role of these tissues as major contributors of plasma lipids during fasting and cold (Haemmerle et al., 2006; Heine et al., 2018; Simcox et al., 2017). Most BAT lipids that were significantly changed with cold exposure were also positively predictive of plasma ACars. Since BAT is known to catabolize lipids for thermogenic fuel, the regression analysis may be detecting tissue that contribute to circulating ACars (liver and iWAT) as well as tissues that are major consumers of ACars (BAT).

**Figure 4:**
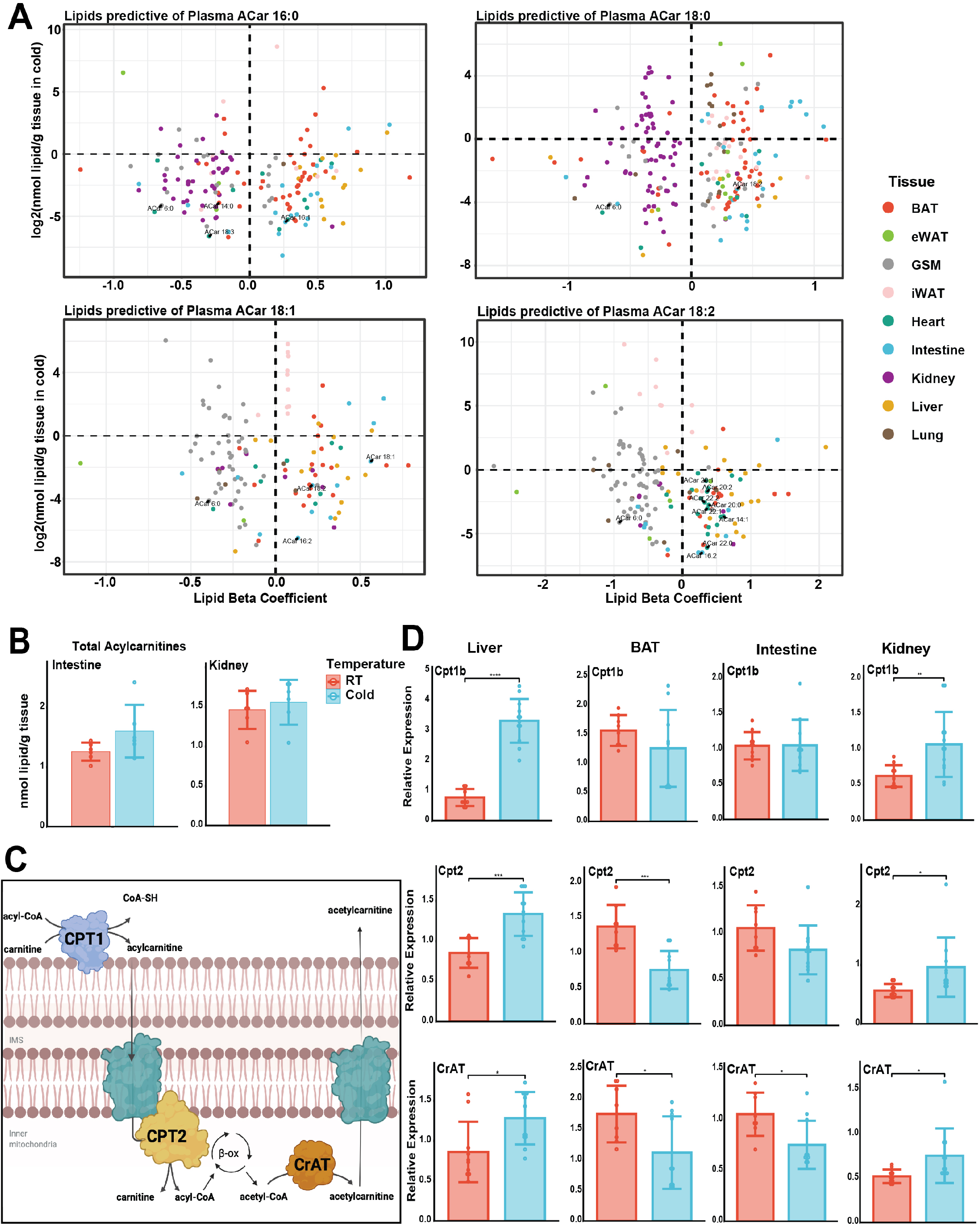
Regression analysis identifies the kidney and intestine as novel regulators of circulating acylcarnitines. A) Regression analysis to predict the tissue sources of major plasma acylcarnitines (ACars). Only lipids significantly predictive (*P*<0.05) are included. ACars are labeled in black text and the tissue of origin of lipid predictors is indicated by color. B) Total ACars in intestine and kidney after 6h at room temperature (RT, 24°C) or cold (Cold, 4°C). C) ACar metabolism showing key enzymes involved in the pathway. D) RT-PCR analysis of ACar metabolism related gene expression in several tissue in RT versus cold mice. Target gene expression was normalized to the housekeeping gene RPS3. Student’s t-test used for all pairwise comparisons; \**P*<0.05, \*\**P*<0.01, \*\*\**P*<0.001, \*\*\*\**P*<0.0001.

We observed that intestine ACars were significant predictors of all major plasma ACars (**Figure 4A**). This was intriguing as there is growing interest in the intestine as an important contributor of circulating lipids, similar to the liver (Han et al., 2021). Total intestine ACars trended higher in cold but only two species, ACar 16:0 and 20:1, were significantly increased (**Figure 4B; Figure S4A**). Though the increase in liver ACars during cold (**Figure 3A**) was more widespread than intestine, contribution to the plasma lipidome does not necessitate intra-tissue lipid changes and it was possible that the intestine trafficked ACars despite not accumulating them. Numerous kidney lipids were negative predictors of plasma ACars despite no changes in total kidney ACars during cold (**Figure 4A-B; Figure S4A**).

Because ACar production is known to be regulated at the transcriptional level, we compared transcriptional changes of genes involved in ACar metabolism between RT and cold in the liver, BAT, intestine, and kidney (**Figure 4C-D**). Expression of all liver transcripts related to ACar metabolism were significantly increased in the cold, supporting higher production of ACar for export to plasma (**Figure 4C; Figure S4B**). BAT uptake and catabolism of circulating ACars was supported by decreases in gene expression of ACar producing enzymes *Cpt1a, Cpt1b*, and *Cpt2*, well as a decrease in *Crat*, which is known to regulate pyruvate dehydrogenase indicating that a decrease in *Crat* would signal the shift from glucose to lipid catabolism (Muoio et al., 2012). These observations align with previous findings, including the increases of *Cpt1b* in the liver with cold even though it was originally characterized as a muscle specific isoform (McGarry et al., 1977; Simcox et al., 2017; Sun et al., 2021). ACar producing enzymes were not significantly changed in intestine during cold, however, *Crat* expression was decreased. This indicates that while production of intestine ACars is not significantly perturbed in cold, there may be a decrease in mitochondrial export leading to net absorption ACars from circulation by intestine. Interestingly, kidney ACar transcripts had identical patterns to that of the liver. These data support a role of increased production and contribution of ACars from the kidney into circulation during cold.

Through regression and complementary gene expression analyses, we were able to confirm the liver and BAT as respective contributors and consumers of circulating ACars. We also identified the intestine and kidney as regulators of circulating ACars, a function not previously known for these tissues. These results provide a proof of principle for computational identification of tissue contribution to the plasma pool and warrant the further investigation of ACar metabolism and function in the intestine and kidney during cold exposure.

### BAT and kidney contribute to the circulating ceramide pool

In addition to ACars, ceramides, a type of sphingolipid, were the most increased lipids in plasma, liver, and BAT during cold exposure (**Figure 2C**). Ceramides are known to communicate cellular identity as well as regulate insulin signaling and fatty acid oxidation (Summers et al., 2019; Tan-Chen et al., 2020). There was a significant increase in total ceramides in plasma and liver but not BAT during cold exposure, though the overall level of ceramides was higher in BAT than plasma or liver regardless of temperature (**Figure 5A; Figure S5A**). When we compared levels of all detected ceramide species in RT and cold in plasma, we observed that the total ceramide increase was primarily driven by Cer d18:1_22:0, Cer d18:1_24:0, and Cer d18:1_24:1 (**Figure 5B; Figure S5B**).

**Figure 5:**
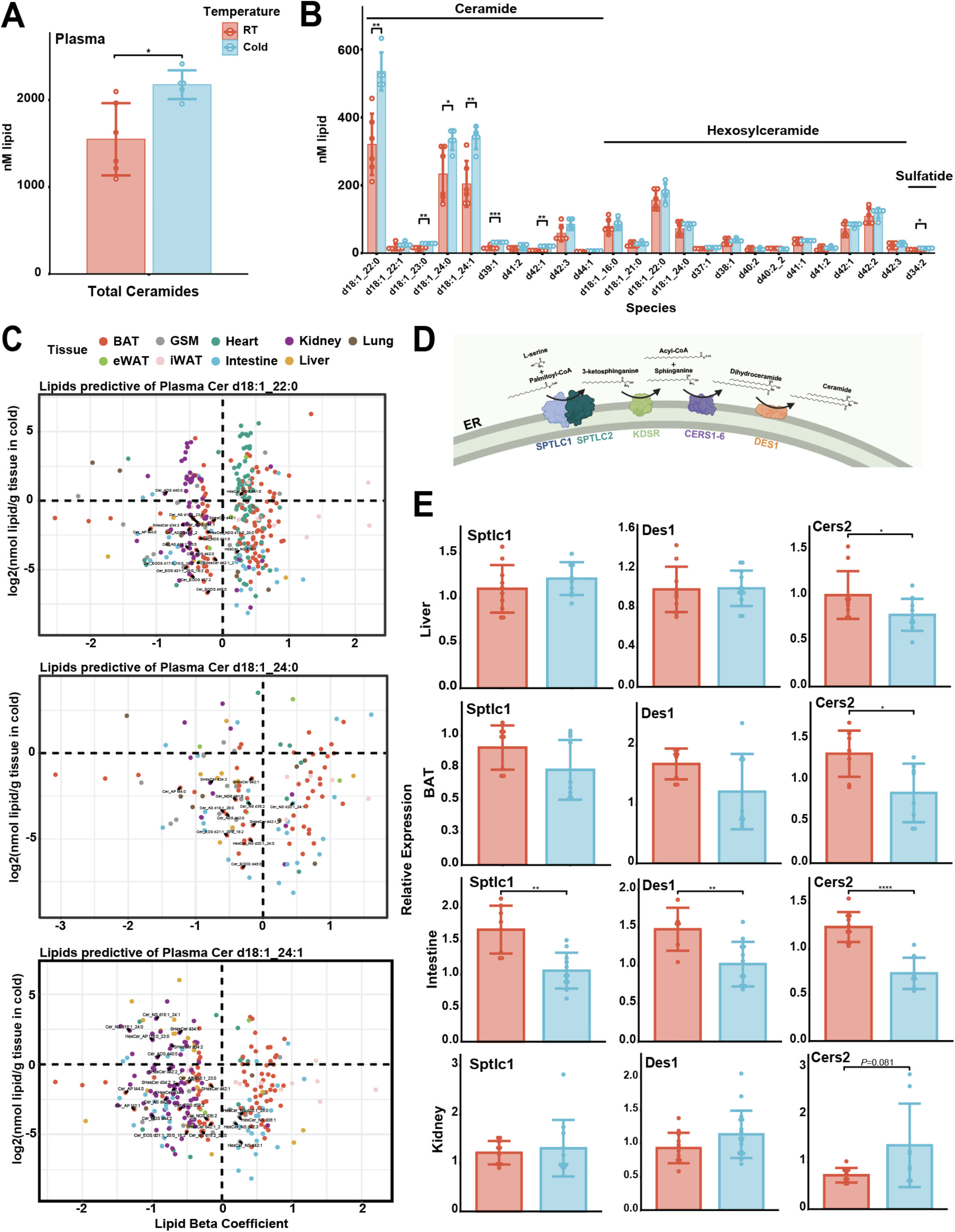
Shifts in ceramide levels occur across multiple tissue during cold exposure. A) Total ceramide abundance in plasma after 6h at room temperature (RT, 24°C) or cold (Cold, 4°C). B) Abundance of all ceramide species detected in plasma at RT or cold. C) Regression analysis to predict the tissue sources of major plasma ceramides (Cers). Only lipids significantly predictive (*P*<0.05) are included. Cers are labeled in black text and the tissue of origin of lipid predictors is indicated by color. D) Cer synthesis pathway showing key enzymes involved in the pathway. E) RT-PCR analysis of Cer metabolism related gene expression in several tissue in RT versus cold mice. Target gene expression was normalized to the housekeeping gene RPS3. Student’s t-test used for all pairwise comparisons; \**P*<0.05, \*\**P*<0.01, \*\*\**P*<0.001, \*\*\*\**P*<0.0001.

To determine which tissue may be contributing to the circulating ceramide pool, we performed regression analysis on the major plasma ceramides Cer d18:1_22:0 (d22:0), Cer d18:1_24:0 (d24:0), and Cer d18:1_24:1 (d24:1; **Figure 5C**). Lipids from BAT dominated all three regressions as both positive and negative predictors of circulating ceramides with BAT ceramides generally negatively predictive. This may be indicative of BAT contribution to circulating ceramides, though total ceramides were unchanged in BAT during cold exposure (**Figure S5A**). Liver lipids were poorly represented as significant predictors of circulating ceramides despite an increase in total liver ceramides during cold (**Figure S5A**). This was unexpected given the observation that the liver is the primary regulator of numerous circulating lipids including triglycerides, acylcarnitines, and cholesterol. Intestinal lipids were predictive of all three major circulating ceramides, though there was no discernable pattern regarding positive or negative predictive value. We further observed variation in the tissue prediction of circulating ceramides based on the ceramide species. Many lipids from the heart were positive predictors of d22:0 but not d24:0 or d24:1. Kidney lipids were predominantly negative predictors of circulating d22:0 and d24:1 but not d24:0. Total intestine, heart, and kidney ceramides were unchanged in the cold (**Figure S5A**). The range in diversity of ceramide species also mirrors these observations with the intestine, plasma, and liver exhibiting similar patterns in Cer species, which could indicate a shared source, whereas the other tissue are more diverse (**Figure S6A**).

We performed gene expression analysis of the major ceramide synthesis genes in BAT, liver, intestine, heart and kidney to further characterize their roles in affecting the circulating ceramide pool because transcriptional regulation of ceramide synthesis enzymes is indicative of ceramide production in prolonged cold exposure (**Figure 5D-E; Figure S6B**) (Chaurasia et al., 2016; Chaurasia et al., 2021). BAT expression of the ceramide synthesis genes *Cers2, Cers5* and *Cers6* was significantly decreased. Expression of the rate limiting protein transcript, *Sptl1*, was also decreased, though not significantly, in the cold. Given that total BAT ceramides were unchanged in the cold yet d22:0 and d24:0 were significantly increased (**Figure S6C**), these data support a role of ceramide uptake and potentially remodeling in the BAT. A robust ceramide program in BAT is further supported by the identification of 70 ceramide species – second to only kidney (81 ceramides) - many of which trended higher in cold (**Figure 2C**).

Heart and liver ceramide transcripts were either unchanged or decreased in cold despite an increase in total liver ceramides (**Figure 5E; Figure S5A & C**). Since the major liver ceramides increased were d22:0, d24:0 and d24:1 and as there are few liver lipids predictive of circulating ceramides, the liver may be a destination of circulating ceramides. Intestinal ceramide transcripts were significantly decreased in cold. This included not only the expression of key synthesis enzymes including *Sptlc1-2, Des1* and several *Cers*, but also sphingomyelinases and ceramidases involved in the salvage pathway and remodeling of ceramides (**Figure 5E**; **Figure S5C**). These data strongly indicate a broad downregulation of ceramide pathways in the intestine and the associated lack of total ceramide changes indicates a role of ceramide uptake by the intestine in the cold. In contrast, kidney *Cers2* expression, which controls the incorporation of 22 and 24 carbon chain lengths into ceramides, was increased in cold (*P*=0.081). Though no other kidney ceramide transcripts were significantly increased in cold, most synthesis and remodeling transcripts trended higher in cold, and this was a unique feature to kidney of all tested tissue. These data indicate that the kidney may play an important role as a contributor to circulating ceramides in cold.

The increase in circulating ceramides and dynamic changes in BAT ceramide species and genes indicates the existence of an important ceramide program during thermogenesis. Unlike ACars, the liver does not seem to be a major contributor to circulating ceramides but may be a consumer. Our data also implicates the kidney as an important contributor to circulating ceramides in cold, whereas the intestine may be involved in uptake. Collectively, these data show a multi-organ ceramide program during thermogenesis that results in increased total ceramides in the circulation. The function of these ceramides remains unknown and is an exciting avenue for further research.

## Discussion

Cold exposure is known to induce maximal lipolysis, shuttling free fatty acids from stored WAT depots to sites of elevated fatty acid oxidation. We took a systemic approach to understand the impact of this rapid FFA flux on various tissues after acute cold exposure of 6 hours. Most tissues were minimally impacted by cold exposure. GSM, intestine, and lung exhibited no significant cold induced lipid changes, while heart and kidney exhibited minor decreases in triglycerides and increases in acylcarnitines. The plasma, liver, and BAT exhibited the most remodeling with cold exposure (**Figure 2C**). The two lipid classes that were primarily increased with cold exposure in the plasma were ACars and Cers. We had previously demonstrated that cold exposure was able to increase plasma ACar through increased FFAs processing to ACars in the liver, and then release into the circulation with uptake in the BAT (Simcox et al., 2017). We were able to recapitulate this multi-tissue processing and uptake through computational assessment of untargeted lipidomics of all nine tissues, demonstrating that the liver was the tissue of origin for plasma acylcarnitines (**Figure 4**). Surprisingly, regression analysis also indicated that the intestine and kidney may have a role in cold remodeling of plasma acylcarnitines. Subsequent RT-PCR analysis of acylcarnitine production enzymes indicated that the kidney increased production of ACars while the intestine may take up ACars. Regression analysis also uncovered a multi-organ ceramide program in thermogenesis (**Figure 5**). In totality these results demonstrate that lipid composition of tissues is dynamically regulated by physiological stresses such as cold exposure, and computational assessment of these lipids can inform on tissues of production and uptake.

We have previously shown that serum and liver ACars concurrently increase throughout the course of acute cold exposure. Using a ^14^C-ACar 16:0 tracer, the fate of circulating ACars during cold was tracked to catabolism in BAT to fuel thermogenesis (Simcox et al., 2017). In the current study, we recapitulated these findings and extended them by profiling not only the liver and plasma ACars, but also the different species present in BAT and their changes in cold (**Figure 2C-D; STable 1**). Further, we found that ACar species with 20+ carbon chains were readily detectable in solid tissue but not in circulation, reflecting differences in the fate of ACar species during cold (**Figure 3A**). ACars require transporters to move across lipid membranes, though there are currently no known plasma membrane importers of ACars and the only characterized acylcarnitine exporter regulates the efflux of short chain and medium chain acylcarnitines (Berardi et al., 1998; Kim et al., 2017; Nakanishi et al., 2001). It is possible that export of ACars cannot occur above a certain acyl length, potentially a clue towards identifying candidate transporters of ACars in the plasma membrane. This would indicate similarities to FFAs with 20+ carbons which, at least in the liver, are only known to be exported in lipoproteins as they are poorly soluble in blood (Bremer and Norum, 1982). Acylcarnitines above 14 carbons travel in the hydrophilic plasma bound to albumin, while short chain acylcarnitines are freely mobile in the circulation to further facilitate quick entry into tissue mitochondria, and this function may be adversely affected if the acyl chain is too long.

We used linear regressions as unbiased computational tools to predict the sources of circulating ACars (**Figure 4A**). We were able to confirm the validity of this method by identifying liver and BAT lipids as significant predictors of circulating ACars. However, we had to use complementary gene expression analysis to elucidate the differing roles of liver and BAT as contributors and consumers, respectively, of plasma ACars. These findings highlight the utility and shortcomings of regressions to identify important predictors, but not their function, when considering the relationship between tissue and plasma lipidomes. This is not unique to lipid-lipid analyses but is common in genetic network analyses as well (Salleh et al., 2017). Our regression and RT-PCR analyses also identified the kidney and intestine as potential contributors and consumers, respectively, of circulating ACars (**Figure 4A; Figure 4C**). Similar to the liver, the kidney is known to take up circulating lipids in the form of FFA during fasting (Hui et al., 2020; Scerbo et al., 2017). Interestingly, kidney lipids were only significant predictors for saturated ACars 16:0 and 18:0, but not unsaturated 18:1 or 18:2, and liver ACars correlated better with unsaturated ACar 18:1 than saturated 18:0 (**Figure 3B**). This raises the possibility that the liver and kidney contribution to plasma ACars is dependent on ACar species, which may be regulated at the level of substrate availability or transport mechanisms of ACars in the respective tissue. It has been previously shown that there is a modest increase of intestinal ACars in 24h fasted rats (Sachan and Ruark, 1985). Our results support these findings as we saw a modest but non-significant increase of intestinal ACars after 6h of cold (**Figure 4B**). Together, our results provide novel insight into the role of kidney and intestine as regulators of circulating ACars during cold. Further investigation into the precise mechanisms and importance of these tissue in lipid-mediated cold adaptation is warranted.

We observed a cold induced rise in several Cers species in the plasma with cold exposure. This observation is striking given that we and others observed that sphingomyelin is the most abundant class of sphingolipid in the plasma followed by ceramides, monohexosylceramides, and dihexosylceramide (**Figure 1C**) (Muralidharan et al., 2021). Regression analysis determined that these Cer species may be coming from the BAT and kidney. Several of the most abundant ceramide species in circulation were also increased in BAT with cold exposure (**Figure 2C**) and there were no HexCers changes, as others have observed with acute treatment of β-adrenergic receptor agonist (Chaurasia et al., 2016). We also established that the liver is a major regulator of plasma Cers, likely through uptake, and that total liver Cers are increased with cold exposure (**Fig 5E**; **Figure S5A**). The functional role of increased total plasma and liver Cers in acute thermoregulation is unknown. A potential explanation may be related to lipoprotein metabolism. In addition to triglyceride uptake and ACar production, clearance of high-density lipoproteins is an important function of the liver in cold (Bartelt et al., 2017). Still poorly understood, reverse cholesterol transport results in the efflux of cholesterol from peripheral tissue into HDL destined for clearance by the liver. As cholesterol and ceramide are known to associate in endosomal pathways, which contribute to cholesterol efflux, a potential explanation for increased liver ceramides may be through clearance of HDL particles which adsorb ceramides that are associated with cholesterol (Goldschmidt-Arzi et al., 2011). Future studies on the mechanisms of ceramide transport through circulation should include assessment of exosome and HDL particles which will inform on the functional role in various physiological stresses.

Cers have been shown to regulate differentiation in brown and beige adipocytes, and adipose tissue-specific knockout of Cer synthesis enzyme *Sptlc2* led to decreased Cers, increased mitochondrial content, increased respiration, and increased thermogenic transcript expression (Chaurasia et al., 2016; Chaurasia et al., 2021; Li et al., 2020; Summers et al., 2019). Conversely, loss of Cer degradation through knockout of acid ceramidase 1 driven by the UCP1 promoter led to increased ceramide levels, decreased mitochondria, decreased respiration, and increased thermogenic transcripts (Chaurasia et al., 2021). This regulation is beyond differentiation capacity since acute treatment of beige adipocytes with a sphingolipid synthesis inhibitor, myriocin, increased cellular respiration, while increasing Cer levels by treating beige adipocytes with C2-ceramide led to decreased respiration (Chaurasia et al., 2016). Beyond regulation of differentiation and respiration, Cer species are known to regulate insulin sensitivity (Chaurasia et al., 2019; Li et al., 2020; Summers et al., 2019). Perhaps in acute cold exposure, where a rapid shift from glucose to lipid catabolism is needed to conserve glucose stores for the brain, Cer induced insulin resistance confers an adaptive advantage. More work is needed to demonstrate how plasma Cers levels are regulated since expression of Cer synthases decreased in BAT and liver with cold exposure (**Figure 5E**), two of the most well characterized tissue in cold exposure. Intriguingly, our data shows that the kidney is an important determinant of circulating ceramides (**Figure 5C & E**) and others have shown that it is a major consumer of circulating lipids (Hui et al., 2020; Scerbo et al., 2017). The role of the kidney in acute thermogenesis and lipid homeostasis is poorly understood and answering these questions could inform on the dynamics of plasma ceramides and their functional role in cold.

One of the major challenges in lipidomics is accurate annotation on global lipidomics sets. The program we used for annotation is LipidAnnotator which has coverage of 58 lipid types including ether and oxidized lipids using the LipidBlast library (Kind et al., 2013). While LipidAnnotator provides substantial coverage, its annotation also includes duplications that highlight the need for library curation and further processing to limit the impact on computational assessment (Koelmel et al., 2020). We used retention time correlation analysis to determine if duplicate annotations were indicative of true replication, which can occur when combining positive and negative mode, or if they represent spectral overlap of distinct species that cannot be distinguished. Other challenges with annotation in global lipidomics include over-annotation, particularly of sphingolipids due to the range and structural similarities numerous species differentiated solely by a double bond or hydroxyl group. Others have observed similar issues with sphingolipid annotation (Hartler et al., 2020; Köfeler et al., 2021). This highlights the need for global lipidomics to be validated by inspection of matched spectra for independent verification, and retention time corrections. Compounding on these difficulties is the heterogeneity of various tissues that contain numerous cell types. Liver in particular, which we found to be an important regulator of circulating ACars and Cers, has been shown to exhibit zonation in cholesterol and glycerolipid synthesis (Ben-Moshe and Itzkovitz, 2019). Future development of single cell technologies including single cell metabolomics will inform on the contribution of each cell type, but further work is needed in the development of mass spectrometry imaging to provide spatial distinction.

The findings in this paper are a systematic assessment of lipid extraction from various tissues, computational imputation of tissue contribution to the plasma lipid pool, and characterization of acute lipid dynamics. We observed that the variability in lipid extraction between methods is different with each tissue and in the end, we utilized single phase extraction for ease of protocol and broad coverage of lipid classes between tissues. Application of optimized global lipidomics to each tissue allowed us to determine that acute cold exposure increases plasma ACars and Cers, and identify tissues that regulate these circulating pools. These studies are an important step to establishing the functional role of plasma lipids and determining the mechanism through which they are transported into the circulation.

## Acknowledgments

The authors would like to thank and acknowledge the contribution of the Simcox laboratory for help with tissue harvest as well as Mae Hurtado-Thiele and Paula Gonzalez for assistance in sample preparation for extraction. The authors would also like to thank the Simcox laboratory for manuscript review including Helaina Von Bank, Edrees Rashan, and Gisela Geoghegan. We would also like to acknowledge the contribution of Greg Barrett-Wilt and Timothy Shriver from the Mass Spectrometry Core Facility in the UW-Madison Biotechnology Center for maintaining the QTOF-LC/MS. Research reported in this publication was supported by the Eunice Kennedy Shriver National Institute of Child Health & Human Development of the National Institutes of Health, the Office of The Director, National Institutes of Health (OD) and the National Cancer Institute (NCI) under Award Number K12HD101368. The content is solely the responsibility of the authors and does not necessarily represent the official views of the National Institutes of Health. This award was supported in part by UW-Madison Comprehensive Diabetes Center Core and the University of Wisconsin - Madison Office of the Vice Chancellor for Research and Graduate Education with funding from the Wisconsin Alumni Research Foundation. The work was also supported in part by startup funds from the University of Wisconsin-Madison School Department of Biochemistry to JS. Other funds that supported this publication include funds from the Diabetes Research Center at Washington University in St. Louis of the National Institutes of Health under award number P30DK020579. B.C. received research support from the National Institutes of Health (DK115824 and DK124326); American Diabetes Association Career Development Award (7-21-JDF-033) and the USDA (2019-67018-29250).

## Author Contributions

RJ, GW, and JS were responsible for conceptualization, data analysis, experimentation, and manuscript preparation. RJ and IO developed and advised on computational analysis pipeline, respectively. RJ, GW, BC, IO, and JS contributed to methods development, editing, revisions, and literature searches.

## STAR Methods

### Resource Availability

All raw LC/MS data will be deposited in MetaboLights. All R code for data processing and curated data for the analysis will be available on Github. All other materials are available upon request to the corresponding author.

### Mouse Husbandry

All animal procedures were approved by the Institutional Animal Care and Use Committee (IACUC) at the University of Wisconsin-Madison. Mice were housed at room temperature (21-23°C) and 40-70% humidity using a 12-hour light/12-hour dark cycle. Mice were fed a standard chow diet (5008 Formulab) and given *ad libitum* access to food and water. Male C57BL6J mice aged 12-14 weeks were used in all experiments. Mice were purchased from Jackson Laboratories.

### Cold Tolerance Test

Beginning at zetgeiber time 3, mice were placed at either 22°C (room temperature) or 4°C and 30% humidity (cold exposure) for 6 hours. For the duration of the cold tolerance test in mice were singly housed with no food or bedding but free access to water. Animals were evaluated hourly for abnormal physiological changes.

### Tissue Collection and Processing

Mice were anesthetized by isoflurane and euthanized by cervical dislocation. Blood samples were collected by cardiac puncture in tubes containing 0.5 mL 129 mM buffered sodium citrate (Covidien) for plasma extraction. Excised tissue was washed in sterile PBS prior to sectioning and snap freezing.

### Reagents and Standards

All reagents were LCMS grade or better. Chloroform, methanol and IPA were purchased from Sigma Aldrich (Cat no. #132950, #1060351000 and #34863, respectively), ethyl acetate from Honeywell (#UN1173) and methyl tert-butyl ether (MTBE) and water from Thermo Fisher Scientific (#E1274 and #51140, respectively). Butylated hydroxytoluene (BHT; Sigma-Aldrich #B1378) was added as an antioxidant to methanol or IPA at a concentration of 0.02% (w/v) unless otherwise stated. Avanti SPLASH Lipidomix (Cat. No. 330707-1EA) and oleoyl-carnitined3 (Cayman Chemical #26578) were used as internal standards (IS).

### Lipid Extractions

For all methods, ten microliters of SPLASH mix and 300 pmol oleoyl-carnitine_d3_ was added per sample as IS, and processing blanks without IS were run for each method. Tissue was homogenized in ceramic 1.4mm bead tubes (Cat. No. 13113-50) using the Qiagen TissueLyzer II (Cat. No. 9244420) for 2-6 cycles using chilled (4°C) blocks (**Supplementary Methods**). Homogenized samples were centrifuged at 4°C for 10 mins at 16,000g to induce phase separation and/or pellet extracted protein prior to organic solvent transfer into new 1.5mL microcentrifuge tubes. Extractions were done on ice using cold solvents. Lipid extracts were dried in a SpeedVac and resuspended in 100% methanol (MeOH) without butylated hydroxytoluene (BHT) for all tissue except iWAT and eWAT, which were resuspended in 100% IPA for better solubility of triglycerides. Samples were stored at −20°C for no more than one week and freeze-thawed <3 times prior to analysis.

#### Folch method

Tissue was homogenized in 2:1 CHCl3:MeOH containing IS at a volume of 100uL per 5mg tissue or 20μL plasma and 150μL water was added to induce phase separation (Folch et al., 1957; Reis et al., 2013). Tubes were inverted several times, centrifuged, and the top aqueous layer was transferred to a new 1.2mL tube. 500μL of 2:1 solution was added to the aqueous layer for re-extraction, the tube was inverted, centrifuged, and the top aqueous layer was aspirated. A glass pipette was used to pool the organic layers in a 1.5mL tube prior to drying.

#### MTBE method

Tissue was homogenized in 300μL of methanol containing IS. 500μL of MTBE was added, followed by 300μL water to induce phase separation. Tubes were inverted to mix, centrifuged, and the top organic layer was extracted into a new 1.5mL tube. Remaining aqueous phase was re-extracted with 500μL MTBE and pooled with the first extract for drying (Matyash et al., 2008).

#### Acidified MTBE method

The MTBE method was repeated with a slight modification. After extracting the initial organic layer, 100μL of MeOH with 1% formic acid (v/v) was used to acidify the aqueous layer prior to re-extraction with 500μL of MTBE (Reis et al., 2013).

#### IPA Method

Tissue was homogenized in 500uL of a 3:1:6 IPA:H_2_O:ethyl acetate solution containing IS (Bielawski et al., 2006; Sarafian et al., 2014). Following homogenization, samples were placed in −20°C for 10 min to precipitate protein, centrifuged, and the entire solvent layer was extracted into a new tube. If tissue particles were still visible, samples were re-spun and supernatant was transferred to a new tube for drying.

### LC/MS Parameters

Extracts were separated on an Agilent 1260 Infinity II UHPLC system using an Acquity BEH C18 column (Waters 186009453, 1.7 μm 2.1 x 100 mm) maintained at 50°C with VanGuard BEH C18 precolumn (Waters 18003975). The chromatography gradient comprised of mobile phase A containing ACN:H_2_O (60:40), 10 mM ammonium formate, and 0.1% formic acid and mobile phase B containing 9:1:90 ACN/H_2_O/IPA, 10 mM ammonium formate, and 0.1% formic acid run at a flow rate of 0.5mL/min. The mobile phase gradient for positive and negative ionization modes began with 15% mobile phase B increased to 30% over 2.40 min, then increased to 48% until 3 min, next to 82% at 13.2 min, then increased to 99% from 13.2-13.8 min and held from 13.8-15.4 min before re-equilibration to 15%, held until 20 min.

The UHPLC system was connected to an Agilent 6546 Q-TOF MS dual AJS ESI mass spectrometer. For positive mode, the gas temperature was kept at 250°C, gas flow at 12 L/min, nebulizer at 35 psig, sheath gas temperature at 300°C, and sheath gas flow at 11 L/min. Negative mode gas temperature and flow were the same while the nebulizer was kept at 30 psig, sheath gas temperature at 375°C, and sheath gas flow at 12 L/min. The VCap voltage was set at 4000V, skimmer at 75V, fragmentor at 190V, and Octopole RF peak at 750V for both ionizations. Samples were injected in a random order and scanned between 100 and 1500 m/z. Reference masses used for positive mode were 121.05 and 922.00 m/z. For negative mode, they were 112.98 and 966.00 m/z. Tandem MS was performed at a fixed collision energy of 25V. The injection volume was 3μL for positive mode and 5μL for negative mode.

### Gene Expression

RNA was isolated from liver, BAT, GSM, heart, kidney, and intestine using TRIzol reagent (Invitrogen). Samples were homogenized with a TissueLyzer II (Qiagen). Reverse transcription was performed with High-Capacity cDNA Reverse Transcription Kit (Thermo Fisher). Quantification of gene expression was performed with PowerUp SYBR Green 2x Master Mix (Thermo Fisher) on an Applied Biosystems QuantStdui 5 Real-Time PCR Syste, 384-well. Relative expression was extrapolated from a standard curve for each primer pair and normalized to expression of the housekeeping gene RPS3.

### Data Processing

Raw data was collected in .d format and MS/MS data was analyzed using Agilent MassHunter Qualitative Analysis and LipidAnnotator for lipid identification (Koelmel et al., 2020). Lipids identified in LipidAnnotator were exported to PCDL format to create individual, comprehensive libraries for each tissue. Identifications for select lipids from different class were checked for accuracy through retention time correlations with lipids of the same class and fragmentation pattern assessment, in addition. Data to compare the lipidome of cold versus room temperature mice was collected in MS1 and imported into Agilent Profinder for lipid identification and peak integration using the tissue specific libraries. Data was exported to .csv files and in-house R scripts were used for normalization to internal standards and starting tissue amount (*R* version 4.0.2).

Lipid annotations were further screened for redundancies through criteria based on analysis of raw chromatograms. The reasons for these redundancies included multiple adducts of the same lipid, slight retention time differences corresponding to the same lipid peak, and in-source fragmentation. As these redundant annotations were also present for lipid standards, we developed additional filtering criteria to obtain high confidence, unique lipid identifications. For positive ionization, individual identifications (ids) for the same lipid were filtered by dropping any duplicates if retention time difference for id <0.10 min or if the lower intensity annotation was <25% of the more abundant id. Cutoffs were determined by evaluating duplicate annotations for internal standards. To determine total unique identifications, positive and negative mode data were compared for identical lipids annotated in both modes and the lower abundance id was dropped. All raw data files have been deposited to Chorus (#) and *R* code is available on Github (URL). Any other files are available upon request.

### Statistics

Statistical analyses of lipidomic data was performed in R using *tidyverse* (Hadley Wickham, 2019). Lipid data was normalized to the appropriate IS and reported in pmol lipid/mg tissue except for plasma, which is in units pmol lipid/mL plasma. Principal component analyses were conducted using the *factoextra* package. Mean ± SD was reported and *P*-values less than 0.05 were considered significant unless otherwise stated. A *n*=6 mice was used for all treatment groups. Mice were considered outliers if lipid values were >2 SD in a given comparison. To determine which lipids were changed in cold for each tissue, the log2(fold change) in cold versus RT was plotted against the −log_10_(t-test *P*-value) for each lipid. *P*-values were corrected for false discovery and considered significant if *q*<0.30 as this was a discovery-based analysis. Pearson correlations were used to determine relationships between lipid species in different tissue after outlier removal.

For regressions, lipid data was log_2_ transformed, mean-centered and scaled to the standard deviation. Each lipid identified in solid tissue was then individually regressed against the plasma lipid of interest with temperature as a covariate using the following model:

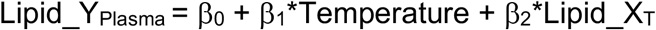

where Lipid_Y_plasma_ was the plasma lipid of interest, β_0_ was the y-intercept (effectively zero due to mean centering), temperature was a binomial where zero was cold and one was RT, and Lipid_X was a given lipid from tissue (T) being tested as a potential predictor. Lipids were considered significantly predictive if both the temperature and lipid terms had a *P*<0.05 as this was indicative of predictive power after accounting for the independent effect of temperature. We also tested regression models with inclusion of an interaction between lipid and temperature, but the term was not significant in our regressions. Further information on packages used throughout the analyses can be found in the **Supplementary Methods**.

## Key Resource Table

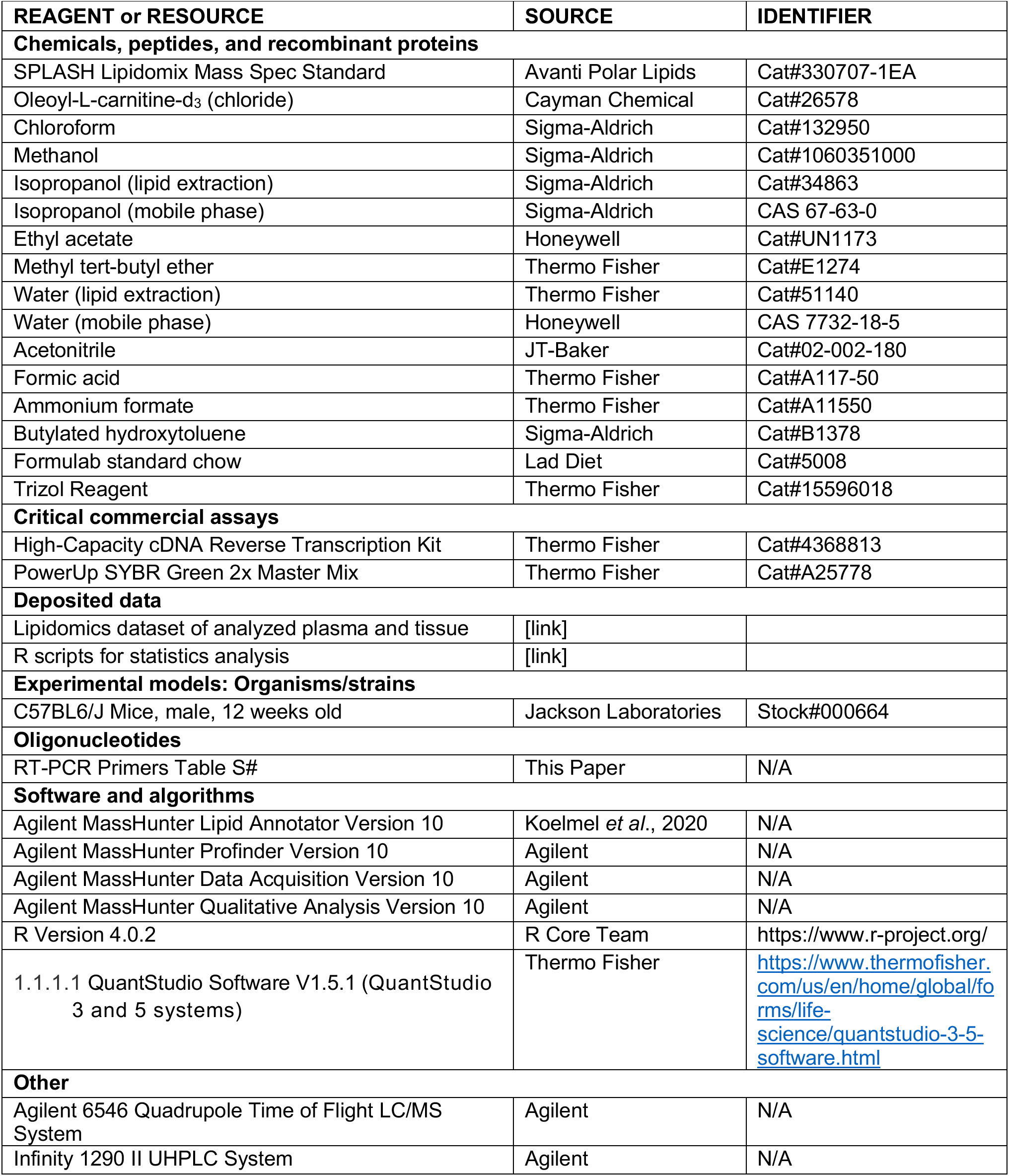

## RT-PCR Primer Table

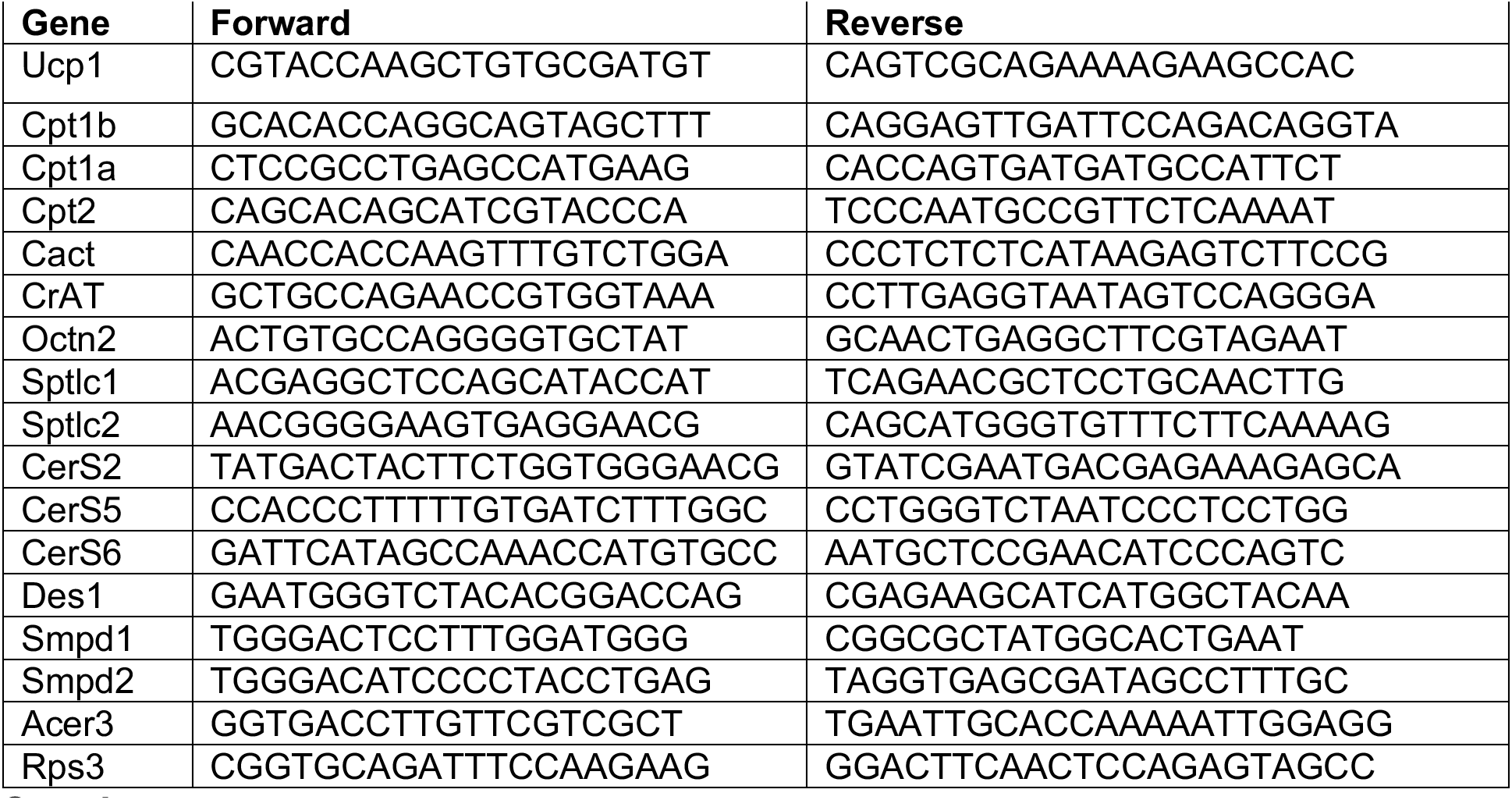

## Supplement

**Supplementary methods.** Details on tissue amount and dilution for lipidomics and further explanation of *R* packages used in analysis.

**Supplemental Figure 1.** A) Breakdown of lipid identification distribution for liver and intestine across the four extraction methods. Lipids were identified in LipidAnnotator following LC/QTOF-MS/MS data collection on pooled plasma from male C57BL6J mice (n=3).

**Supplemental Figure 2.** A) Comparison of total lipid abundances for the major lipid classes measured in the tissue of male C57BL6J mice housed at room temperature (RT; 24°C) or cold (4°C) for 6h. n=6 for each condition. B) Volcano plot showing individual lipid changes across multiple tissue in RT versus cold. Data corrected for multiple comparison using FDR and *q*<0.30 considered significant and plotted in red (lipids increased in cold) or blue (lipid decreased in cold). Student’s t-test used for all pairwise comparisons; \**P*<0.05 \*\**P*<0.01 \*\*\**P*<0.001.

**Supplemental Figure 3.** A) Pearson correlation analysis between plasma, liver and brown adipose tissue (BAT) for ACar 14:0, ACar 16:0 and 18:2. B) Venn diagram showing the dominant acyl chain by lipid abundance in plasma, liver and BAT for mice kept at room temperature (24°C; RT) or cold (4°C) for 6h. C) BAT and inguinal white adipose tissue total ceramide breakdown by sphingoid base (d18:0 or d18:1) at RT versus cold. Student’s t-test used for all pairwise comparisons; \**P*<0.05.

**Supplemental Figure 4.** A) Comparison of changes in all acylcarnitine (ACar) species detected in intestine and kidney, respectively, for mice kept at room temperature (24°C; RT) or cold (4°C) for 6h. B) RT-PCR showing gene expression of proteins involved in ACar metabolism for tissue at RT versus cold. Student’s t-test used for all pairwise comparisons; \**P*<0.05 \*\**P*<0.01 \*\*\**P*<0.001.

**Supplemental Figure 5.** A) Comparison of total ceramide abundance in tissue of mice kept at room temperature (24°C; RT) or cold (4°C) for 6h. B) Representative MS/MS spectra confirming identification of major ceramides d18:1_22:0, d18:1_24:0 and d18:1_24:1 from liver raw mass spec data. Spectra from LipidAnnotator viewer with library in blue and actual data in red. Ceramide figures from LipidMaps (lipidmaps.org). C) RT-PCR showing gene expression of proteins involved in ceramide metabolism in tissue of RT versus cold mice. D) Comparison of all brown adipose tissue ceramide species at RT and cold. Student’s t-test used for all pairwise comparisons; \**P*<0.05 \*\**P*<0.01 \*\*\**P*<0.001.

**Supplemental Table 1.** Lipid species increased in plasma, liver and brown adipose tissue as well as overlap from Venn Diagram in Figure 2D.

