## Supplement to Figures 1-5 for "Systematic assessment of lipid profiles for the discovery of tissue contributors to the circulating lipid pool in cold exposure"

**Supplementary methods**

The following tissue amounts and dilutions were used for analysis:

| Tissue | Target Weight (+/- 2mg) | Pos dilution | Inj volume Pos | Neg dilution | Inj volume neg |
|---|---|---|---|---|---|
| **Liver** | 25 | 45x | 3uL | undiluted | 5uL |
| **BAT** | 12 | 70x | 3uL | undiluted | 5uL |
| **iWAT** | 15 | 150x | 3uL | undiluted | 5uL |
| **eWAT** | 15 | 150x | 3uL | undiluted | 5uL |
| **Skeletal Muscle** | 15 | 30x | 3uL | undiluted | 5uL |
| **Lung** | 25 | 35x | 3uL | undiluted | 5uL |
| **Heart** | 15 | 20x | 3uL | undiluted | 5uL |
| **Intestine** | 25 | 45x | 3uL | undiluted | 5uL |
| **Kidney** | 25 | 30x | 3uL | undiluted | 5uL |
| **Plasma** | 100uL | 20x | 3uL | undiluted | 5uL |

*Inj = injection; iWAT = inguinal white adipose tissue; eWAT = epididymal WAT

The *reshape* package was used for data wrangling (1). The packages were used *ggpubr* and *ggrepel* packages was used for bar graph aesthetics and labeling points in volcano plots (2,3). *rstatix* was used for pairwise comparisons (4). *ggsci* and *RColorBrewer* were used to make color palettes (5,6). *VennDiagram* was used to construct Venn Diagrams (7). The *heatmaply* package was used to visualize lipid class breakdown across extractions and different tissue (8). *Cairo* was used to output pdf files for further editing (9).

**Package citations**


1. H. Wickham. Reshaping data with the reshape package. Journal of Statistical Software, 21(12), 2007.
2. Alboukadel Kassambara (2020). ggpubr: 'ggplot2' Based Publication Ready Plots. R package version 0.4.0. https://CRAN.R-project.org/package=ggpubr
3. Kamil Slowikowski (2021). ggrepel: Automatically Position Non-Overlapping Text Labels with 'ggplot2'. R package version 0.9.1. https://CRAN.R-project.org/package=ggrepel
4. Alboukadel Kassambara (2021). rstatix: Pipe-Friendly Framework for Basic Statistical Tests. R package version 0.7.0. https://CRAN.R-project.org/package=rstatix
5. Nan Xiao (2018). ggsci: Scientific Journal and Sci-Fi Themed Color Palettes for 'ggplot2'. R package version 2.9. https://CRAN.R-project.org/package=ggsci
6. Erich Neuwirth (2014). RColorBrewer: ColorBrewer Palettes. R package version 1.1-2. https://CRAN.R-project.org/package=RColorBrewer
7. Hanbo Chen (2018). VennDiagram: Generate High-Resolution Venn and Euler Plots. R package version 1.6.20. https://CRAN.R-project.org/package=VennDiagram
8. Tal Galili, Alan O'Callaghan, Jonathan Sidi, Carson Sievert; heatmaply: an R package for creating interactive cluster heatmaps for online publishing, Bioinformatics, , btx657, https://doi.org/10.1093/bioinformatics/btx657
9. Simon Urbanek and Jeffrey Horner (2020). Cairo: R Graphics Device using Cairo Graphics Library for Creating High-Quality Bitmap (PNG, JPEG, TIFF), Vector (PDF, SVG, PostScript) and Display (X11 and Win32) Output. R package version 1.5-12.2. https://CRAN.R-project.org/package=Cairo

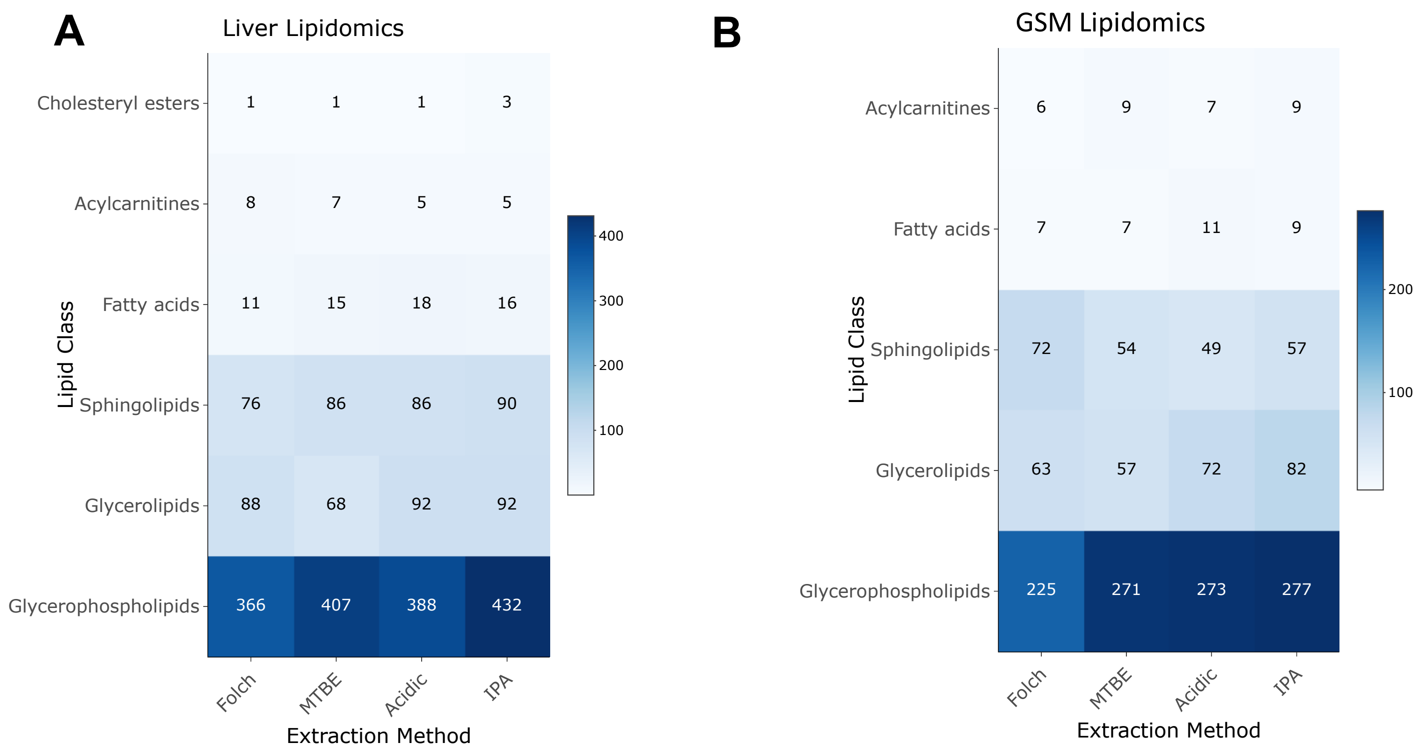


**Supplementary Figure 1.** A) Breakdown of lipid identification distribution for liver and B) intestine across the four extraction methods. Lipids were identified in LipidAnnotator following LC/QTOF-MS/MS data collection on pooled plasma from male C57BL6J mice (n=3).

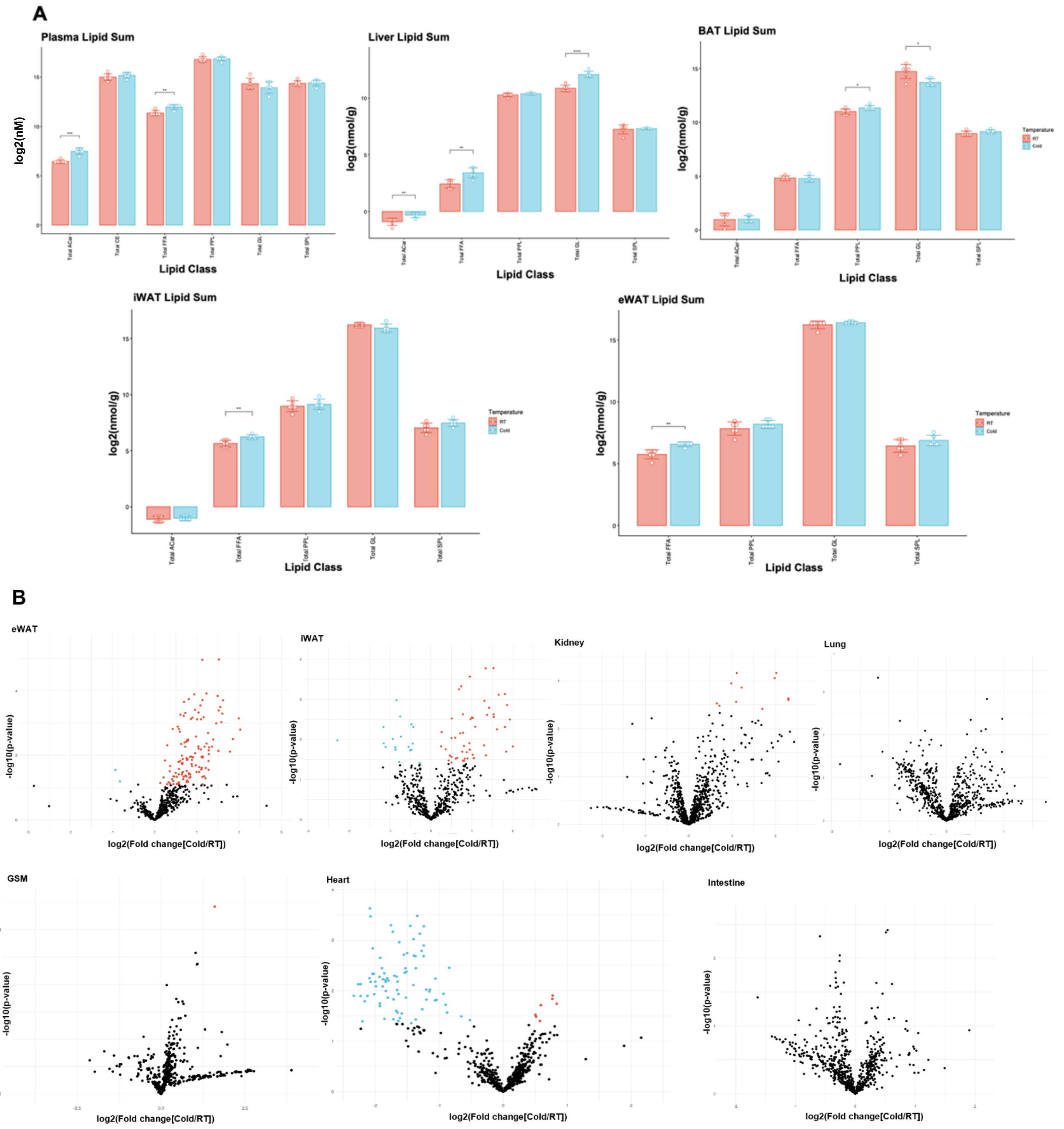


**Supplemental Figure 2.** A) Comparison of total lipid abundances for the major lipid classes measured in the tissue of male C57BL6J mice housed at room temperature (RT; 24°C) or cold (4°C) for 6h. n=6 for each condition. B) Volcano plot showing individual lipid changes across multiple tissue in RT versus cold. Data corrected for multiple comparison using FDR and *q*<0.30 considered significant and plotted in red (lipids increased in cold) or blue (lipid decreased in cold). Student's t-test used for all pairwise comparisons; **P*<0.05 ***P*<0.01 ****P*<0.001.

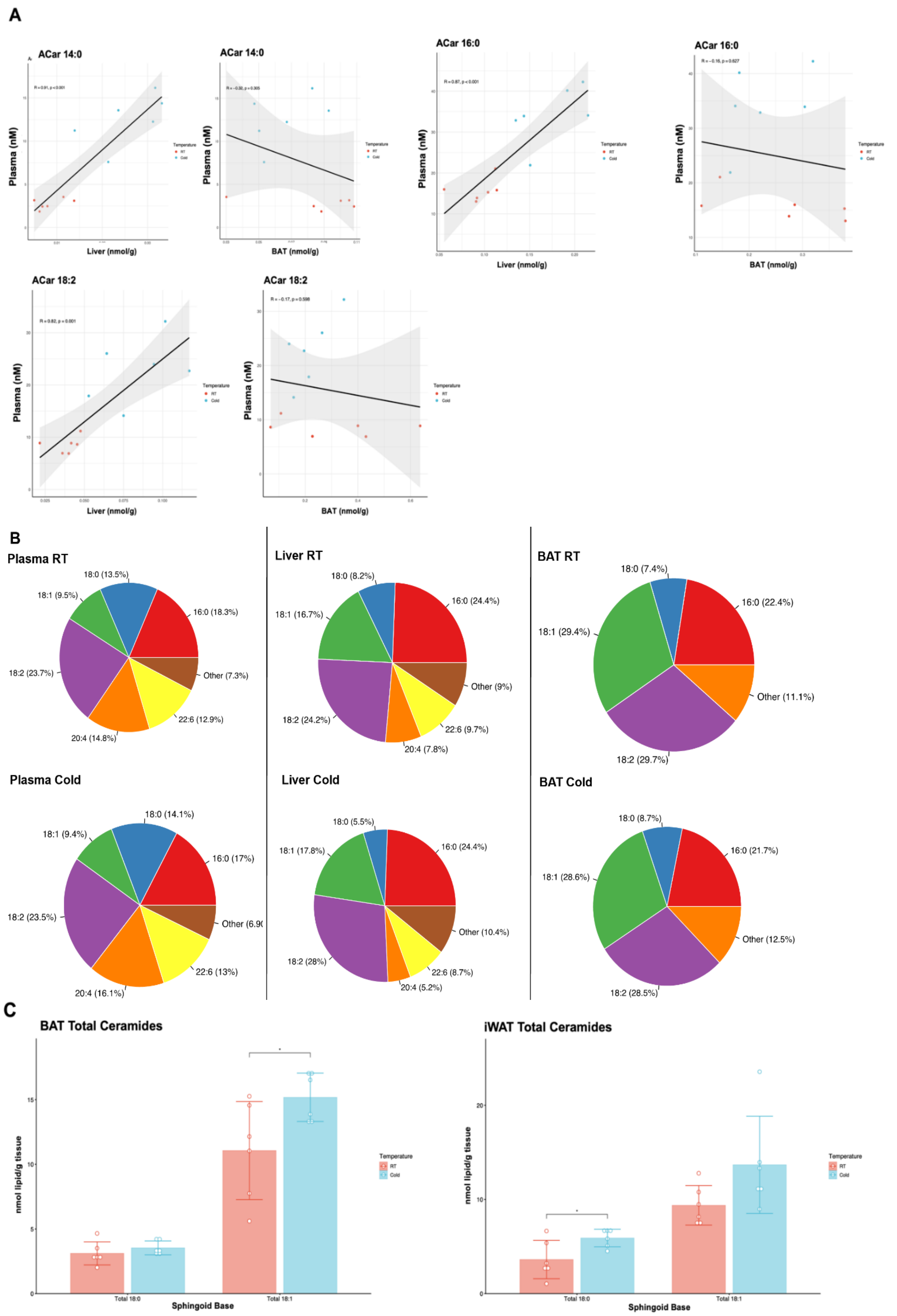


**Supplemental Figure 3.** A) Pearson correlation analysis between plasma, liver and brown adipose tissue (BAT) for ACar 14:0, ACar 16:0 and 18:2. B) Pie charts showing the dominant acyl chain by lipid abundance in plasma, liver and BAT for mice kept at room temperature (24°C; RT) or cold (4°C) for 6h. C) BAT and inguinal white adipose tissue total ceramide breakdown by sphingoid base (d18:0 or d18:1) at RT versus cold. Student's t-test used for all pairwise comparisons; **P*<0.05.

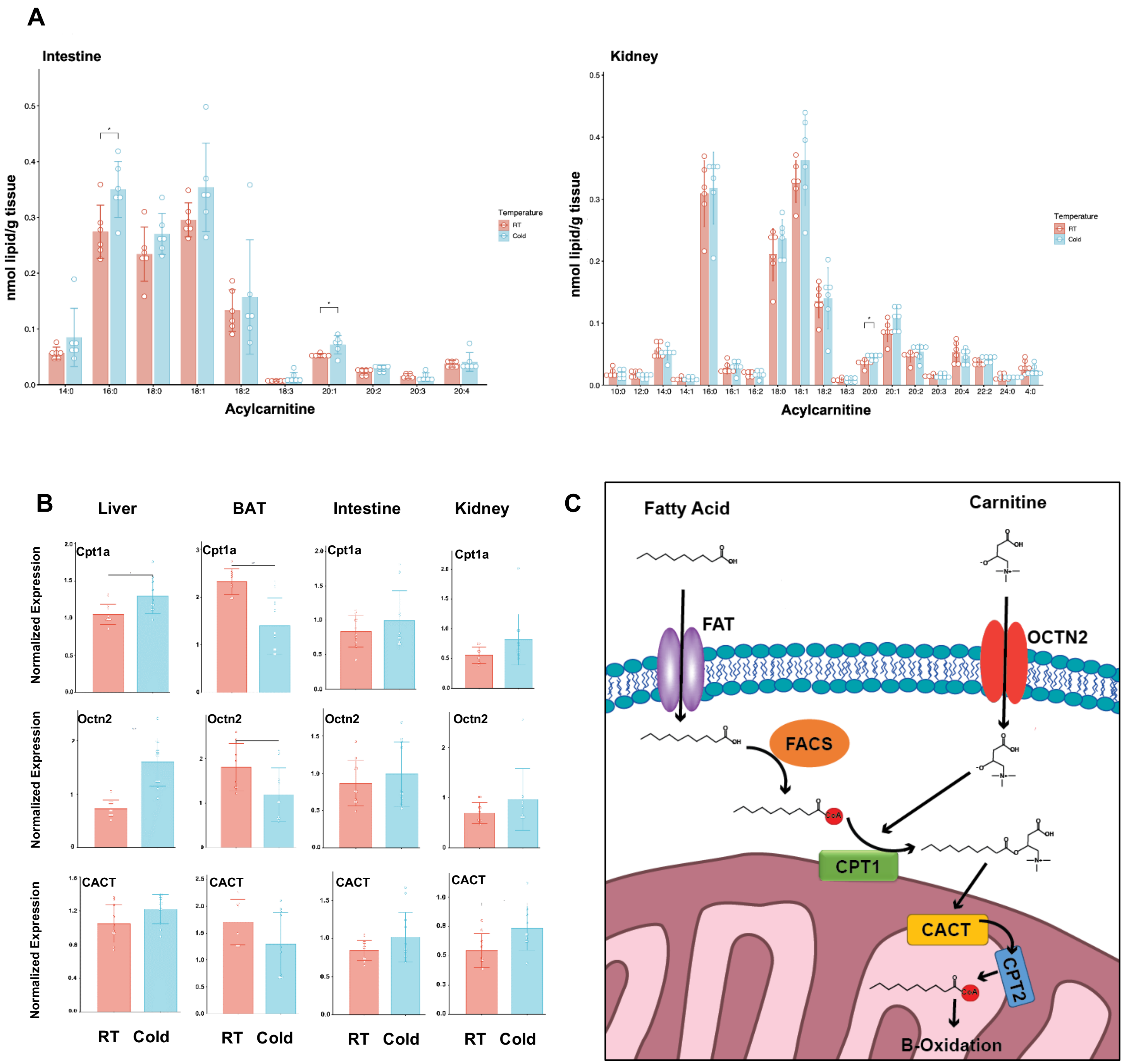


**Supplemental Figure 4.** A) Comparison of changes in all acylcarnitine (ACar) species detected in intestine and kidney, respectively, for mice kept at room temperature (24°C; RT) or cold (4°C) for 6h. B) RT-PCR showing gene expression of proteins involved in ACar metabolism for tissue at RT versus cold. C) Schematic of acylcarnitine processing proteins. Student's t-test used for all pairwise comparisons; **P*<0.05 ***P*<0.01 ****P*<0.001.

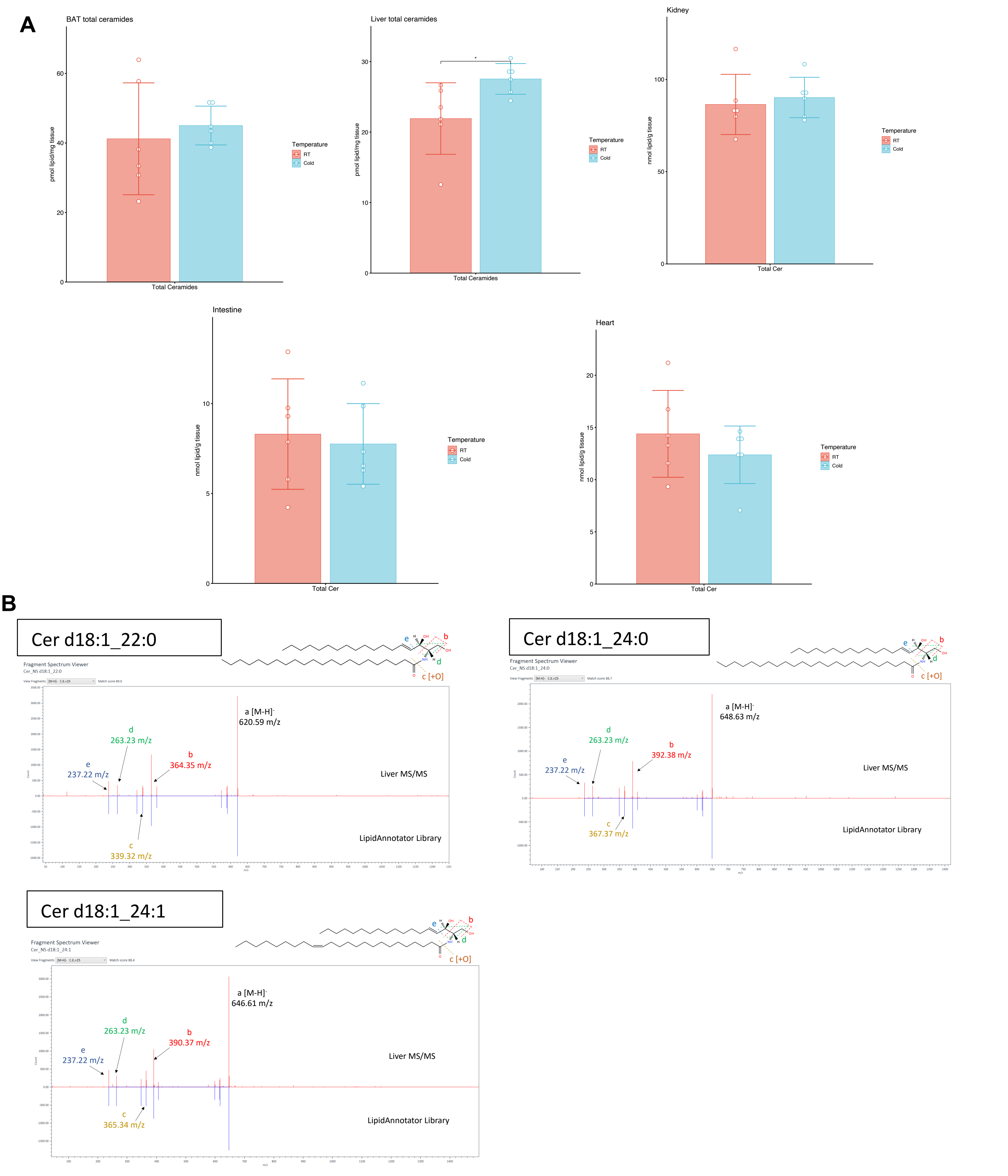


**Supplemental Figure 5.** A) Comparison of total ceramide abundance in tissue of mice kept at room temperature (24°C; RT) or cold (4°C) for 6h. B) Representative MS/MS spectra confirming identification of major ceramides d18:1_22:0, d18:1_24:0 and d18:1_24:1 from liver raw mass spec data. Spectra from LipidAnnotator viewer with library in blue and actual data in red. Ceramide figures from LipidMaps (lipidmaps.org).

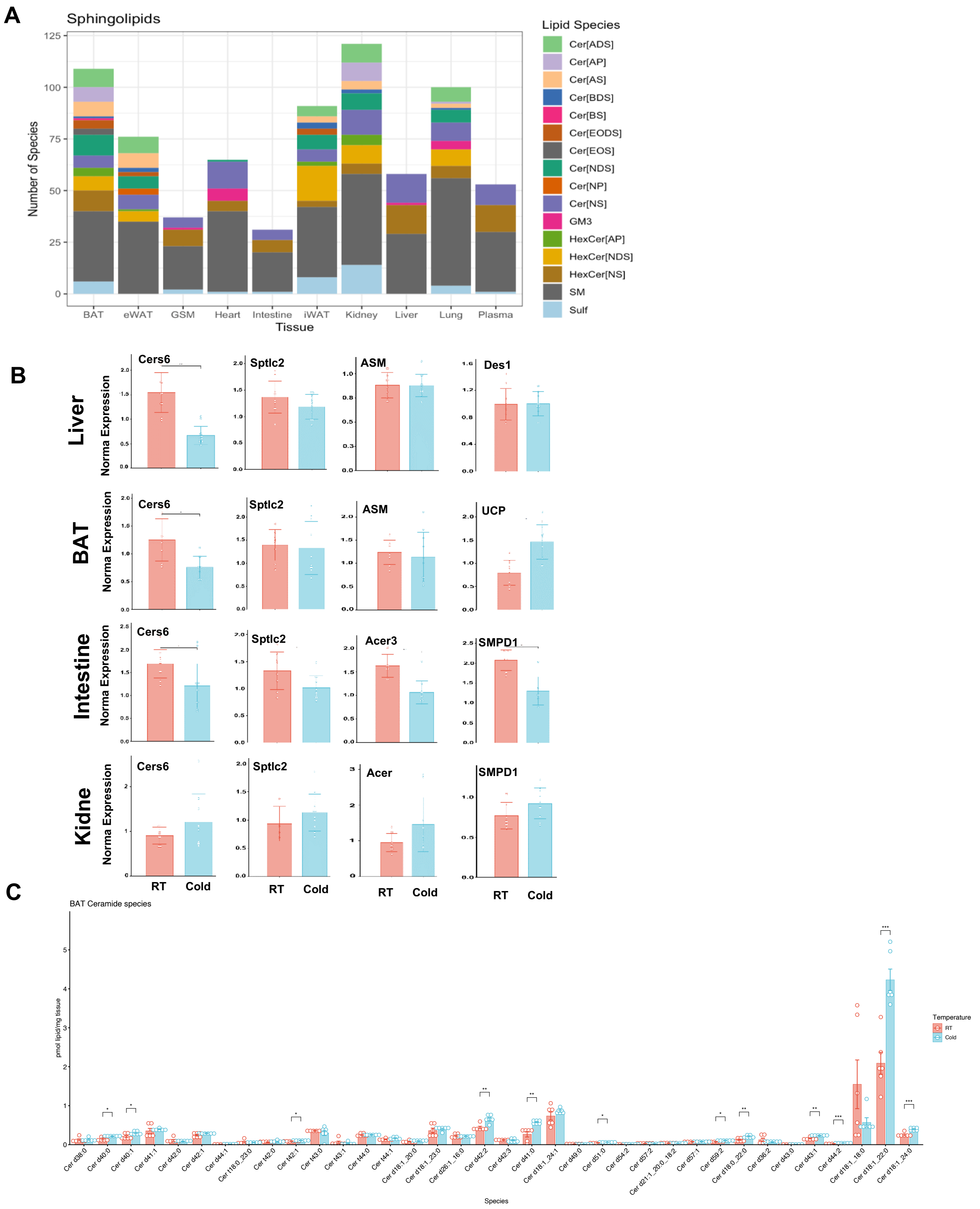


**Supplemental Figure 6.** A) Comparison of ceramides species between different tissues. B) RT-PCR showing gene expression of proteins involved in ceramide metabolism in tissue of RT versus cold mice. C) Comparison of all brown adipose tissue ceramide species at RT and cold. Student's t-test used for all pairwise comparisons; *$P$<0.05 **$P$<0.01 ***$P$<0.001.

**Supplemental Table 1.** Lipid species increased in plasma, liver and brown adipose tissue as well as overlap from Venn Diagram in Figure 2D.

| Plasma_only | Liver_only | BAT_only | Liver_plasma | BAT_plasma | BAT_liver | all_shared |
|---|---|---|---|---|---|---|
| ACar 12:0 | ACar 20:4 | ACar 6:0 | ACar 14:0 | Cer_NS d42:1 | Cer_NS d18:1_22:0 | ACar 18:0 |
| ACar 12:1 | BMP 18:2_22:6 | Cer_ADS d40:0 | ACar 16:0 | FA 20:2 | Cer_NS d18:1_24:0 | Cer_NS d18:1_23:0 |
| ACar 14:1 | Cer_NS d17:1_22:0 | Cer_ADS d40:1 | ACar 18:1 | NA | LPC 14:0/0:0 | Cer_NS d18:1_24:1 |
| ACar 14:2 | Cer_NS d18:1_20:0 | Cer_AP t42:1 | ACar 18:2 | NA | LPC 20:1/0:0 | FA 20:3 |
| ACar 16:1 | Cer_NS d18:2_22:0 | Cer_AS d42:2 | FA 18:1 | NA | LPE 18:0 | FA 22:4 |
| ACar 16:2 | Cer_NS d18:2_24:1 | Cer_BDS d41:0 | FA 18:2 | NA | LPG 18:1 | PC 34:4 |
| CE 20:4 | Cer_NS d36:2 | Cer_EODS d51:0 | FA 18:3 | NA | LPG 18:2 | NA |
| Cer_NS d39:1 | Cer_NS d40:1 | Cer_EOS d59:2 | FA 20:1 | NA | LPI 18:1 | NA |
| EtherPE 18:1e_22:4 | Cer_NS d42:4 | Cer_NDS d18:0_22:0 | FAHFA 18:1_18:0 | NA | PC 14:0_18:2 | NA |
| FA 16:0 | Cer_NS d43:2 | Cer_NDS d41:1 | FAHFA 18:1_20:3 | NA | PC 14:0_20:4 | NA |
| FA 17:1 | EtherPC 18:0e_20:4 | Cer_NDS d42:0 | FAHFA 18:2_18:1 | NA | PC 15:0_18:2 | NA |
| FA 22:5 | EtherPC 18:1e_20:4 | Cer_NDS d42:1 | FAHFA 18:2_20:4 | NA | PC 16:0_18:3_2 | NA |
| FAHFA 16:0_18:2 | EtherPE 18:0e_18:2 | Cer_NDS d43:1 | LPI 20:4 | NA | PC 16:1_18:1 | NA |
| PC 16:2_18:2 | EtherPE 18:0e_20:4 | Cer_NDS d43:1_2 | PC 16:1_18:2 | NA | PC 16:1_18:3 | NA |
| PC 32:2 | FA 18:3_2 | Cer_NDS d44:2 | TG 52:8 | NA | PC 18:0_18:1 | NA |
| SHexCer d34:2 | FAHFA 22:6_22:5 | CL 64:3 | NA | NA | PC 18:0_22:5 | NA |
| NA | HBMP 62:13 | CL 66:1 | NA | NA | PC 18:1_18:1 | NA |
| NA | LPA 16:0 | CL 66:2 | NA | NA | PC 18:2_18:3 | NA |
| NA | LPC 15:0/0:0 | CL 66:3 | NA | NA | PC 33:1 | NA |
| NA | LPC 16:1/0:0 | CL 66:5 | NA | NA | PC 40:4 | NA |
| NA | LPC 17:1/0:0 | CL 68:2 | NA | NA | PE 19:0_20:4 | NA |
| NA | LPC 18:2/0:0 | CL 68:3 | NA | NA | PE 42:9 | NA |
| NA | LPC 18:2/0:0_2 | CL 68:4 | NA | NA | PG 18:0_18:2 | NA |
| NA | LPC 18:3/0:0 | CL 68:5 | NA | NA | PI 18:0_18:1 | NA |
| NA | LPC 20:0/0:0 | CL 68:5_2 | NA | NA | PI 18:0_18:2 | NA |
| NA | LPC 20:2/0:0 | CL 68:6 | NA | NA | SM d32:1 | NA |
| NA | LPC 20:5/0:0 | CL 70:3 | NA | NA | NA | NA |
| NA | LPC 22:0/0:0 | CL 70:4 | NA | NA | NA | NA |
| NA | LPC 22:1/0:0 | CL 70:5 | NA | NA | NA | NA |
| NA | LPC 22:5/0:0 | CL 70:6 | NA | NA | NA | NA |
| NA | LPC 22:6/0:0_2 | CL 70:7 | NA | NA | NA | NA |
| NA | LPC 24:0/0:0 | CL 70:8_2 | NA | NA | NA | NA |
| NA | LPE 17:0 | CL 72:10 | NA | NA | NA | NA |
| NA | LPE 18:2 | CL 72:3 | NA | NA | NA | NA |
| NA | LPE 18:2_2 | CL 72:4 | NA | NA | NA | NA |
| NA | LPE 19:0 | CL 72:5 | NA | NA | NA | NA |
| NA | LPE 20:0 | CL 72:6 | NA | NA | NA | NA |
| NA | LPE 20:3 | CL 72:6_2 | NA | NA | NA | NA |

| | | | | | | |
|---|---|---|---|---|---|---|
| NA | LPE 20:4 | CL 72:7 | NA | NA | NA | NA |
| NA | LPE 20:4_2 | CL 72:8 | NA | NA | NA | NA |
| NA | LPE 22:6 | CL 72:9 | NA | NA | NA | NA |
| NA | LPE 22:6_2 | CL 74:11_2 | NA | NA | NA | NA |
| NA | LPG 22:6_2 | CL 74:7 | NA | NA | NA | NA |
| NA | LPI 16:0 | CL 74:8 | NA | NA | NA | NA |
| NA | LPI 18:0 | CL 74:8_2 | NA | NA | NA | NA |
| NA | LPI 18:2 | CL 74:9 | NA | NA | NA | NA |
| NA | LPI 19:0 | CL 74:9_2 | NA | NA | NA | NA |
| NA | LPI 20:0 | CL 76:11_2 | NA | NA | NA | NA |
| NA | LPI 20:3 | CL 76:12_2 | NA | NA | NA | NA |
| NA | LPI 22:5 | CL 76:13 | NA | NA | NA | NA |
| NA | LPI 22:6 | EtherPC 16:0e_16:0 | NA | NA | NA | NA |
| NA | LPS 16:0 | EtherPC 16:0e_18:2 | NA | NA | NA | NA |
| NA | LPS 22:6 | EtherPC 16:1e_16:0 | NA | NA | NA | NA |
| NA | PA 16:0_18:2_2 | EtherPC 16:2e_20:4 | NA | NA | NA | NA |
| NA | PC 14:0_18:3 | EtherPC 18:1e_18:2 | NA | NA | NA | NA |
| NA | PC 14:0_22:6 | EtherPC 38:5e | NA | NA | NA | NA |
| NA | PC 16:0_16:1 | GlcADG 18:0_20:1 | NA | NA | NA | NA |
| NA | PC 16:0_17:0 | GlcADG 38:0 | NA | NA | NA | NA |
| NA | PC 16:0_19:1 | GlcADG 43:1 | NA | NA | NA | NA |
| NA | PC 16:1_16:1 | HexCer_AP t18:0_22:0 | NA | NA | NA | NA |
| NA | PC 16:1_22:6 | LPC 19:0/0:0 | NA | NA | NA | NA |
| NA | PC 17:2_18:2 | LPC 20:4/0:0_2 | NA | NA | NA | NA |
| NA | PC 18:0_22:4 | LPE 18:1_2 | NA | NA | NA | NA |
| NA | PC 18:0_22:6 | LPG 16:0 | NA | NA | NA | NA |
| NA | PC 18:0_22:6_2 | LPI 18:2_2 | NA | NA | NA | NA |
| NA | PC 18:1_22:5 | PC 14:0_14:0 | NA | NA | NA | NA |
| NA | PC 18:2_18:2 | PC 14:0_16:0 | NA | NA | NA | NA |
| NA | PC 18:2_18:3_2 | PC 14:0_20:5 | NA | NA | NA | NA |
| NA | PC 18:2_18:4 | PC 16:0_16:0 | NA | NA | NA | NA |
| NA | PC 18:3_18:3 | PC 16:0_17:1 | NA | NA | NA | NA |
| NA | PC 19:0_20:4 | PC 16:0_18:0 | NA | NA | NA | NA |
| NA | PC 19:0_22:6 | PC 16:0_18:1 | NA | NA | NA | NA |
| NA | PC 31:0 | PC 16:0_18:2 | NA | NA | NA | NA |
| NA | PC 35:4 | PC 16:0_18:3 | NA | NA | NA | NA |
| NA | PC 35:7 | PC 16:0_20:3 | NA | NA | NA | NA |
| NA | PC 36:6 | PC 16:0_20:4 | NA | NA | NA | NA |
| NA | PE 16:0_18:0 | PC 16:0_20:5 | NA | NA | NA | NA |
| NA | PE 16:0_18:1 | PC 17:0_18:1 | NA | NA | NA | NA |
| NA | PE 16:0_18:3 | PC 17:0_18:2 | NA | NA | NA | NA |
| NA | PE 16:1_18:2 | PC 17:0_20:4 | NA | NA | NA | NA |
| NA | PE 16:1_18:3 | PC 17:1_18:2 | NA | NA | NA | NA |

| | | | | | | |
|---|---|---|---|---|---|---|
| NA | PE 16:1_20:4 | PC 18:0_18:0 | NA | NA | NA | NA |
| NA | PE 18:0_18:1 | PC 18:0_18:2 | NA | NA | NA | NA |
| NA | PE 18:1_18:2 | PC 18:0_20:1 | NA | NA | NA | NA |
| NA | PE 18:2_18:2 | PC 18:0_20:2 | NA | NA | NA | NA |
| NA | PE 18:2_18:3 | PC 18:0_20:3 | NA | NA | NA | NA |
| NA | PE 18:2_18:3_2 | PC 18:0_20:3_2 | NA | NA | NA | NA |
| NA | PE 18:2_20:4 | PC 18:0_20:4 | NA | NA | NA | NA |
| NA | PE 18:3_18:3 | PC 18:0_20:5 | NA | NA | NA | NA |
| NA | PE 19:0_18:2_2 | PC 18:1_18:2 | NA | NA | NA | NA |
| NA | PE 20:0_20:3 | PC 18:1_18:3 | NA | NA | NA | NA |
| NA | PG 16:1_18:2_2 | PC 18:1_20:1 | NA | NA | NA | NA |
| NA | PG 18:1_18:2 | PC 18:1_20:3 | NA | NA | NA | NA |
| NA | PG 18:1_20:4_2 | PC 18:2_20:2 | NA | NA | NA | NA |
| NA | PG 18:2_18:2 | PC 18:2_20:4 | NA | NA | NA | NA |
| NA | PG 18:2_18:3 | PC 18:2_20:5 | NA | NA | NA | NA |
| NA | PG 18:2_18:3_2 | PC 18:2_22:6 | NA | NA | NA | NA |
| NA | PG 18:2_22:6 | PC 19:0_18:2 | NA | NA | NA | NA |
| NA | PG 20:2_22:6 | PC 19:0_18:2_2 | NA | NA | NA | NA |
| NA | PI 18:0_22:6 | PC 20:4_22:6 | NA | NA | NA | NA |
| NA | SM d21:1_19:1 | PC 36:0 | NA | NA | NA | NA |
| NA | SM d36:1 | PC 36:2 | NA | NA | NA | NA |
| NA | SM d42:1_2 | PC 37:7 | NA | NA | NA | NA |
| NA | TG 12:0_14:0_22:4 | PC 38:5 | NA | NA | NA | NA |
| NA | TG 12:0_16:0_18:2 | PC 39:5 | NA | NA | NA | NA |
| NA | TG 12:0_16:1_18:2 | PE 16:0_22:6 | NA | NA | NA | NA |
| NA | TG 12:0_18:2_18:2 | PE 19:0_20:4_2 | NA | NA | NA | NA |
| NA | TG 12:0_18:2_18:3 | PE 20:3_22:6 | NA | NA | NA | NA |
| NA | TG 14:0_16:0_18:1 | PE 40:9 | NA | NA | NA | NA |
| NA | TG 14:0_16:0_18:2 | PG 14:0_18:2 | NA | NA | NA | NA |
| NA | TG 15:0_16:0_18:1 | PG 18:0_22:6_2 | NA | NA | NA | NA |
| NA | TG 15:0_18:1_18:2 | PG 18:2_20:4 | NA | NA | NA | NA |
| NA | TG 15:0_18:2_18:2 | PG 20:1_18:2 | NA | NA | NA | NA |
| NA | TG 16:0_16:0_16:0 | PG 38:6 | NA | NA | NA | NA |
| NA | TG 16:0_16:0_18:0 | PI 16:0_18:2 | NA | NA | NA | NA |
| NA | TG 16:0_16:0_18:1 | PI 16:1_18:1 | NA | NA | NA | NA |
| NA | TG 16:0_16:0_18:2 | PI 16:1_22:4 | NA | NA | NA | NA |
| NA | TG 16:0_16:0_22:6 | PI 18:1_18:2 | NA | NA | NA | NA |
| NA | TG 16:0_16:1_18:2 | PI 18:1_20:3 | NA | NA | NA | NA |
| NA | TG 16:0_16:1_20:5 | PI 18:1_20:4 | NA | NA | NA | NA |
| NA | TG 16:0_16:1_22:6 | PI 18:2_20:4 | NA | NA | NA | NA |
| NA | TG 16:0_16:2_18:2 | PI 36:2 | NA | NA | NA | NA |
| NA | TG 16:0_16:3_18:2 | PI 36:4 | NA | NA | NA | NA |
| NA | TG 16:0_17:0_18:0 | PI 38:7 | NA | NA | NA | NA |
| NA | TG 16:0_17:1_18:1 | PI 40:3 | NA | NA | NA | NA |
| NA | TG 16:0_18:0_18:0 | SM d37:1 | NA | NA | NA | NA |

| | | | | | | |
|---|---|---|---|---|---|---|
| NA | TG 16:0_18:0_18:1 | SM d40:1 | NA | NA | NA | NA |
| NA | TG 16:0_18:0_20:4 | SM d40:2 | NA | NA | NA | NA |
| NA | TG 16:0_18:1_18:1 | SM d41:1 | NA | NA | NA | NA |
| NA | TG 16:0_18:1_18:2 | SM d41:2 | NA | NA | NA | NA |
| NA | TG 16:0_18:1_20:0_2 | SM d42:3 | NA | NA | NA | NA |
| NA | TG 16:0_18:1_20:4 | NA | NA | NA | NA | NA |
| NA | TG 16:0_18:1_22:6 | NA | NA | NA | NA | NA |
| NA | TG 16:0_18:2_18:2 | NA | NA | NA | NA | NA |
| NA | TG 16:0_18:2_18:3 | NA | NA | NA | NA | NA |
| NA | TG 16:0_18:2_18:4 | NA | NA | NA | NA | NA |
| NA | TG 16:0_18:2_20:4 | NA | NA | NA | NA | NA |
| NA | TG 16:0_18:2_20:5 | NA | NA | NA | NA | NA |
| NA | TG 16:0_18:2_22:5 | NA | NA | NA | NA | NA |
| NA | TG 16:0_18:2_22:6 | NA | NA | NA | NA | NA |
| NA | TG 16:0_18:3_22:6 | NA | NA | NA | NA | NA |
| NA | TG 16:0_20:4_22:6 | NA | NA | NA | NA | NA |
| NA | TG 16:0_20:5_22:6 | NA | NA | NA | NA | NA |
| NA | TG 16:0_22:5_22:7 | NA | NA | NA | NA | NA |
| NA | TG 16:0_22:6_22:6 | NA | NA | NA | NA | NA |
| NA | TG 16:1_16:2_18:2 | NA | NA | NA | NA | NA |
| NA | TG 16:1_18:2_18:2 | NA | NA | NA | NA | NA |
| NA | TG 16:1_18:2_18:3 | NA | NA | NA | NA | NA |
| NA | TG 16:1_18:2_20:5 | NA | NA | NA | NA | NA |
| NA | TG 16:1_18:2_22:6 | NA | NA | NA | NA | NA |
| NA | TG 16:1_20:5_22:6 | NA | NA | NA | NA | NA |
| NA | TG 16:2_18:2_18:3 | NA | NA | NA | NA | NA |
| NA | TG 16:2_18:2_20:5 | NA | NA | NA | NA | NA |
| NA | TG 16:3_18:2_18:2 | NA | NA | NA | NA | NA |
| NA | TG 17:0_18:1_18:2 | NA | NA | NA | NA | NA |
| NA | TG 18:0_18:1_22:6 | NA | NA | NA | NA | NA |
| NA | TG 18:1_18:1_18:1 | NA | NA | NA | NA | NA |
| NA | TG 18:1_18:1_18:2 | NA | NA | NA | NA | NA |
| NA | TG 18:1_18:1_22:6 | NA | NA | NA | NA | NA |
| NA | TG 18:1_18:2_18:2 | NA | NA | NA | NA | NA |
| NA | TG 18:1_18:2_20:0 | NA | NA | NA | NA | NA |
| NA | TG 18:1_18:2_20:1 | NA | NA | NA | NA | NA |
| NA | TG 18:1_18:2_20:5 | NA | NA | NA | NA | NA |
| NA | TG 18:1_18:2_22:0 | NA | NA | NA | NA | NA |
| NA | TG 18:1_18:2_22:5 | NA | NA | NA | NA | NA |
| NA | TG 18:1_18:2_22:6 | NA | NA | NA | NA | NA |
| NA | TG 18:2_18:2_18:2 | NA | NA | NA | NA | NA |
| NA | TG 18:2_18:2_18:3 | NA | NA | NA | NA | NA |
| NA | TG 18:2_18:2_18:3_2 | NA | NA | NA | NA | NA |
| NA | TG 18:2_18:2_18:4 | NA | NA | NA | NA | NA |

| | | | | | | |
|---|---|---|---|---|---|---|
| NA | TG 18:2_18:2_20:5 | NA | NA | NA | NA | NA |
| NA | TG 18:2_18:2_22:5 | NA | NA | NA | NA | NA |
| NA | TG 18:2_18:2_22:6 | NA | NA | NA | NA | NA |
| NA | TG 18:2_18:3_18:3 | NA | NA | NA | NA | NA |
| NA | TG 18:2_18:3_18:4 | NA | NA | NA | NA | NA |
| NA | TG 18:2_18:3_20:5 | NA | NA | NA | NA | NA |
| NA | TG 18:2_18:3_22:6 | NA | NA | NA | NA | NA |
| NA | TG 18:2_20:4_22:6 | NA | NA | NA | NA | NA |
| NA | TG 18:2_20:5_22:6 | NA | NA | NA | NA | NA |
| NA | TG 18:2_22:6_22:6 | NA | NA | NA | NA | NA |
| NA | TG 42:2 | NA | NA | NA | NA | NA |
| NA | TG 44:2 | NA | NA | NA | NA | NA |
| NA | TG 46:3 | NA | NA | NA | NA | NA |
| NA | TG 48:3 | NA | NA | NA | NA | NA |
| NA | TG 48:4 | NA | NA | NA | NA | NA |
| NA | TG 48:5 | NA | NA | NA | NA | NA |
| NA | TG 50:1 | NA | NA | NA | NA | NA |
| NA | TG 50:3 | NA | NA | NA | NA | NA |
| NA | TG 50:5 | NA | NA | NA | NA | NA |
| NA | TG 50:6 | NA | NA | NA | NA | NA |
| NA | TG 50:6_2 | NA | NA | NA | NA | NA |
| NA | TG 51:3 | NA | NA | NA | NA | NA |
| NA | TG 51:4 | NA | NA | NA | NA | NA |
| NA | TG 52:2 | NA | NA | NA | NA | NA |
| NA | TG 52:3 | NA | NA | NA | NA | NA |
| NA | TG 52:5 | NA | NA | NA | NA | NA |
| NA | TG 52:5_2 | NA | NA | NA | NA | NA |
| NA | TG 52:6 | NA | NA | NA | NA | NA |
| NA | TG 52:6_2 | NA | NA | NA | NA | NA |
| NA | TG 52:7 | NA | NA | NA | NA | NA |
| NA | TG 52:7_2 | NA | NA | NA | NA | NA |
| NA | TG 53:2 | NA | NA | NA | NA | NA |
| NA | TG 53:3 | NA | NA | NA | NA | NA |
| NA | TG 54:1 | NA | NA | NA | NA | NA |
| NA | TG 54:2 | NA | NA | NA | NA | NA |
| NA | TG 54:3 | NA | NA | NA | NA | NA |
| NA | TG 54:4 | NA | NA | NA | NA | NA |
| NA | TG 54:5 | NA | NA | NA | NA | NA |
| NA | TG 54:6 | NA | NA | NA | NA | NA |
| NA | TG 54:6_2 | NA | NA | NA | NA | NA |
| NA | TG 54:7 | NA | NA | NA | NA | NA |
| NA | TG 54:7_2 | NA | NA | NA | NA | NA |
| NA | TG 54:7_3 | NA | NA | NA | NA | NA |
| NA | TG 54:8 | NA | NA | NA | NA | NA |
| NA | TG 54:8_3 | NA | NA | NA | NA | NA |

| | | | | | | |
|---|---|---|---|---|---|---|
| NA | TG 54:8_4 | NA | NA | NA | NA | NA |
| NA | TG 54:9 | NA | NA | NA | NA | NA |
| NA | TG 54:9_2 | NA | NA | NA | NA | NA |
| NA | TG 56:10_2 | NA | NA | NA | NA | NA |
| NA | TG 56:11 | NA | NA | NA | NA | NA |
| NA | TG 56:2 | NA | NA | NA | NA | NA |
| NA | TG 56:3 | NA | NA | NA | NA | NA |
| NA | TG 56:5 | NA | NA | NA | NA | NA |
| NA | TG 56:6 | NA | NA | NA | NA | NA |
| NA | TG 56:7 | NA | NA | NA | NA | NA |
| NA | TG 56:7_2 | NA | NA | NA | NA | NA |
| NA | TG 56:8 | NA | NA | NA | NA | NA |
| NA | TG 56:8_2 | NA | NA | NA | NA | NA |
| NA | TG 56:9 | NA | NA | NA | NA | NA |
| NA | TG 56:9_2 | NA | NA | NA | NA | NA |
| NA | TG 58:10 | NA | NA | NA | NA | NA |
| NA | TG 58:11 | NA | NA | NA | NA | NA |
| NA | TG 58:11_2 | NA | NA | NA | NA | NA |
| NA | TG 58:11_3 | NA | NA | NA | NA | NA |
| NA | TG 58:12 | NA | NA | NA | NA | NA |
| NA | TG 58:8 | NA | NA | NA | NA | NA |
| NA | TG 58:8_2 | NA | NA | NA | NA | NA |
| NA | TG 58:9 | NA | NA | NA | NA | NA |
| NA | TG 58:9_2 | NA | NA | NA | NA | NA |
| NA | TG 60:11 | NA | NA | NA | NA | NA |
| NA | TG 60:12 | NA | NA | NA | NA | NA |
| NA | TG 62:14 | NA | NA | NA | NA | NA |
